# Cytoplasmic accumulation of FUS triggers early behavioral alterations linked to cortical neuronal hyperactivity and defects in inhibitory synapses

**DOI:** 10.1101/2020.06.09.141556

**Authors:** Jelena Scekic-Zahirovic, Inmaculada Sanjuan-Ruiz, Vanessa Kan, Salim Megat, Pierre De Rossi, Stéphane Dieterlé, Raphaelle Cassel, Pascal Kessler, Diana Wiesner, Laura Tzeplaeff, Valérie Demais, Hans-Peter Muller, Gina Picchiarelli, Nibha Mishra, Sylvie Dirrig-Grosch, Jan Kassubek, Volker Rasche, Albert Ludolph, Anne-Laurence Boutillier, Magdalini Polymenidou, Clotilde Lagier-Tourenne, Sabine Liebscher, Luc Dupuis

## Abstract

Gene mutations causing cytoplasmic mislocalization of the RNA-binding protein FUS, lead to severe forms of amyotrophic lateral sclerosis (ALS). Cytoplasmic accumulation of FUS is also observed in other diseases, with unknown consequences. Here, we show that cytoplasmic mislocalization of FUS drives behavioral abnormalities in knock-in mice, including locomotor hyperactivity and alterations in social interactions, in the absence of widespread neuronal loss. Mechanistically, we identified a profound increase in neuronal activity in the frontal cortex of *Fus* knock-in mice *in vivo*. Importantly, RNAseq analysis suggested involvement of defects in inhibitory neurons, that was confirmed by ultrastructural and morphological defects of inhibitory synapses and increased synaptosomal levels of mRNAs involved in inhibitory neurotransmission. Thus, cytoplasmic FUS triggers inhibitory synaptic deficits, leading to increased neuronal activity and behavioral phenotypes. FUS mislocalization may trigger deleterious phenotypes beyond motor neuron impairment in ALS, but also in other neurodegenerative diseases with FUS mislocalization.

## Introduction

Amyotrophic lateral sclerosis (ALS) is the major adult motor neuron disease, with onset usually in the 6^th^ decade of life and death due to respiratory insufficiency and progressive paralysis usually 3 to 5 years after onset of motor symptoms^1-3^. Mutations in the Fused in Sarcoma gene (*FUS*), encoding an RNA-binding protein from the FET family^4,5^, are associated with the most severe forms of ALS^6,7^, clinically presenting with a very early onset and rapid disease progression^8,9^. ALS associated mutations in *FUS* are clustered in the C-terminal region of the FUS protein that includes the atypical PY nuclear localization sequence, and is required for protein entry into the nucleus^6,7,10-12^. The severity of the disease correlates with the degree of impairment of FUS nuclear import^11,12^, and the most severe cases of ALS known to date are indeed caused by mutations leading to the complete truncation of the PY-NLS^8,9^.

A number of clinical and pathological evidence suggest that FUS mislocalization to cytoplasm and subsequent aggregation could be relevant beyond the few ALS-FUS cases. First, *FUS* mutations, although rare in non ALS cases, have been found in cases with frontotemporal dementia, either isolated^13,14^ or as an initial presentation of ALS-FTD^15,16^, as well as in patients with initial chorea^17^, mental retardation^18^, psychosis or dementia^19^ and essential tremor^20^. In the absence of *FUS* mutations, FUS mislocalization^21^ or aggregation^22,23^ were found to be widespread in sporadic ALS. FUS pathology also defines a subset of cases with FTD (FTD-FUS) with prominent atrophy of the caudate putamen^24-26^, concomitant pathology of other FET proteins TAF15 and EWSR1^12,27-30^ and frequent psychiatric symptoms^28^. FUS aggregates have also been observed in spinocerebellar ataxia and Huntington’s disease^31,32^. While FUS mislocalization appears to be a common feature in neurodegenerative diseases, its pathological consequences have not been thoroughly studied beyond motor neuron degeneration.

Neurons with FUS pathology show decreased levels of FUS in the nucleus, that might compromise a number of processes dependent upon proper FUS levels such as transcription and splicing regulation or DNA damage repair^4^. Interestingly, loss of FUS alters the splicing of multiple mRNAs relevant to neuronal function^33,34^, such as *MAPT*, encoding the TAU protein, and alters the stability of mRNAs encoding relevant synaptic proteins such as GluA1 and SynGAP1^35-39^. However, loss of nuclear FUS levels is very efficiently compensated for by autoregulatory mechanisms as well as by other FET proteins. Loss of nuclear FUS remains indeed limited as opposed to loss of nuclear TDP-43 observed in TDP-43 pathology^40^. Indeed, heterozygous *Fus* knock-in mice, which carry one mutant allele leading to cytoplasmic, and not nuclear localization of FUS, only show marginal loss of nuclear FUS due to compensatory overexpression^10,41^. Beyond nuclear loss of function, accumulation of cytoplasmic FUS was found to be a critical event in ALS-FUS in multiple studies in mouse models. For instance, cytoplasmic FUS is necessary to cause motor neuron degeneration in *FUS*-ALS^10,41-46^ as heterozygous *Fus* knock-in mouse models develop mild, late onset muscle weakness and motor neuron degeneration, but not haploinsufficient *Fus* knock-out mice^10,41,46^. To date, there are few studies investigating whether the accumulation of cytoplasmic FUS might lead to phenotypes beyond motor neuron disease. Interestingly, FUS is also found at synaptic and dendritic sites^38,47-51^, and Sahadevan, Hembach et al (co-submitted manuscript) identify synaptic mRNA targets for FUS that are critical for synaptic formation, function and maintenance.

Here, we show that a partial cytoplasmic mislocalization of FUS in heterozygous *Fus* knock-in mice is sufficient to drive a panel of behavioral abnormalities, including locomotor hyperactivity or alterations in social interactions, which preceded motor neuron degeneration. This was accompanied by ventricle enlargement in the absence of widespread neuronal cell loss. Mechanistically, we could identify a profound increase in neuronal activity in the frontal cortex of *Fus* knock-in mice *in vivo*. Furthermore, we observed impaired expression of multiple genes related to neuronal function in the frontal cortex throughout adulthood. Genes differentially regulated were selectively related to inhibitory neurons and we observed ultrastructural and morphological defects of inhibitory synapses accompanied by increased levels of *Fus, Nrxn1* and *Gabra1* mRNAs in synaptosomes of heterozygous *Fus* knock-in mice. Thus, FUS cytoplasmic enrichment is sufficient to trigger inhibitory synaptic deficits leading to increased neuronal activity and behavioral phenotypes. These findings suggest that FUS mislocalization could trigger deleterious phenotypes beyond motor neurons that could be relevant for both ALS-FUS but also other neurodegenerative diseases with FUS mislocalization.

## Results

### Spontaneous locomotor hyperactivity in *Fus*^ΔNLS/+^ mice

Since FUS mislocalization and aggregation are observed in patients with various neurodegenerative diseases, we hypothesized that partial FUS cytoplasmic mislocalization in *Fus*^*ΔNLS/*+^ mice could be sufficient to cause a number of behavioral phenotypes. Two independent cohorts of mice were analyzed at 4 months of age, before the appearance of motor impairment^41^ and 10 months of age. Evaluation of basal motor activity in a familiar environment showed significantly increased locomotor activity in *Fus*^*ΔNLS/*+^ mice over the 3 consecutive days of observation (**Figure 1a-b**). Interestingly, this hyperactivity was observed through the entire night in 4 months old *Fus*^*ΔNLS/*+^ mice (**Figure 1a**), but only during late night hours in older *Fus*^*ΔNLS/*+^ mice (**Figure 1b**). Hyperactivity was not caused by reduced anxiety since *Fus*^*ΔNLS/*+^ mice showed similar preference for peripheral quadrants over central quadrants as *Fus*^+*/*+^ mice in the open field test (**Figure S1a**). Ambulatory distance, duration of ambulation and mean speed of 10-months-old *Fus*^*ΔNLS/*+^ mice were similar to wild type littermates in the open field indicating the absence of hyperactivity in the novel environment (**Figure S1a-b**). To further confirm the lack of anxiety-related phenotype in *Fus*^*ΔNLS/*+^ mice we used the dark/light box test, based on the preference of mice for dark compartments over illuminated places. In this test, 10 months old *Fus*^*ΔNLS/*+^ mice and *Fus*^+*/*+^ mice showed similar latency to enter, similar frequency of transitions and similar time duration to explore illuminated compartment (**Figure S1c**). Thus, *Fus*^*ΔNLS/*+^ mice are hyperactive, but not anxious.

**Figure 1:**
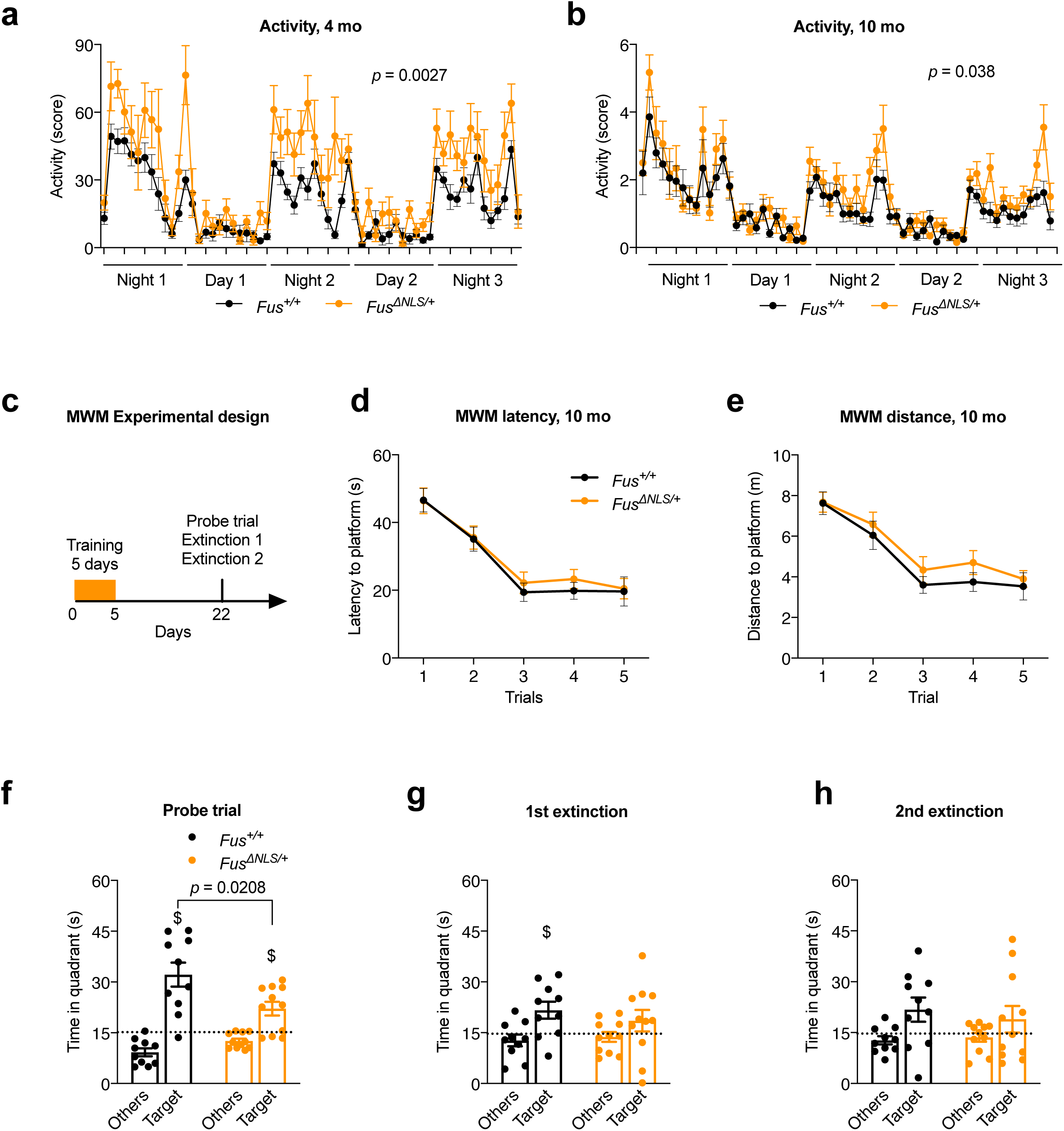
*Fus*^*ΔNLS*/+^ mice display increased nocturnal spontaneous locomotor activity and cognitive defects. (**a-b**) Line graphs represent mice home cage activity – actimetry over three consecutive days at 4 months (**a**) and 10 months (**b**) of *Fus*^+/+^ (black) and *Fus*^*ΔNLS*/+^ (orange) mice. N=10-11 per genotype at 4 months and N=14-15 per genotype for 10 months. p=0.0027 at 4 months and p=0.038 at 10 months for genotype effect (see Table S1 for statistical tests). Data are presented as mean +/- SEM values of activity score per hour. (**c**) Schematic illustration of the Morris water maze (MWM) experimental strategy (paradigm). Mice were subjected to a five-day training period and tested for spatial memory retention in a probe trial (60 seconds) 18 days after the last acquisition. The probe trial was then followed by two extinction tests, performed at 2 hours intervals. (**d-e**) Line graphs represent latency (in seconds) (**d**) and total distance swam (in meters) (**e**) to find the hidden platform during acquisition of 10 months old *Fus*^+/+^ (black) and *Fus*^*ΔNLS*/+^ (orange) mice. Both genotypes improved similarly their performance between day 1 and 5. N=10-11 per genotype. Data are presented as mean +/- SEM values of four trials per day of training. Repeated measures Two-way ANOVA with days and genotype as variables. No significant effect of genotype is observed. (**f**) Bar graphs represent the time spent in the target quadrant (Target) and the average of the time spent in other three quadrants (Others) during probe trial. Dashed line shows chance level (15 seconds per quadrant; i.e. 25%). Both genotypes were significantly above random but *Fus*^*ΔNLS*/+^ mice performed significantly worse than *Fus*^+/+^ littermates. $, p<0.01, Student t-test for comparison to a chance level, Target quadrant: p=0.0008 for *Fus*^+/+^ and p=0.006 for *Fus*^*ΔNLS*/+^. Genotype comparison was made using One-way ANOVA; F(1,19)=6.33, p=0.0208 (**g-h**) Bar graphs represent the time spent in quadrants (Target vs Others) during the first (**g**) and the second (**h**) extinction test. $, p<0.05 vs chance levels. One-way ANOVA for genotype effect (F(1,19)=0.56, p=0.46) (**g**), (F(1,19)=0.27, p=0.60) (**h**) and Student t-test for comparison to a chance level (Target quadrant: p=0.025 for *Fus*^+/+^ and p=0.22 for *Fus*^*ΔNLS*/+^) (**g**), (Target quadrant: p=0.08 for *Fus*^+/+^ and p=0.09 for *Fus*^*ΔNLS*/+^) (**h**).

### Executive dysfunction in *Fus*^ΔNLS/+^ mice

To explore the possibility that behavioral phenotypes of *Fus*^*ΔNLS/*+^ mice included executive dysfunction, we tested spatial reference memory in the Morris water maze. This task requires the hippocampal function, at least during acquisition and memory formation, but relies on a proper cortico-hippocampal dialog for longer retention times or remote memory retrieval (**Figure 1c**)^52^. At 10 months of age, *Fus*^*ΔNLS/*+^ mice performed similarly to their *Fus*^+*/*+^ littermates regarding distance travelled and latency to find hidden platform over training days (**Figure 1d-e**). We then performed a probe trial 18 days after the last training and observed that, although both genotypes searched significantly in the Target quadrant compared to random, *Fus*^*ΔNLS/*+^ mice displayed a significantly decreased performance to retrieve memory at this time point (**Figure 1f**). Furthermore, *Fus*^*ΔNLS/*+^ mice extinguished their previous memory much faster than wild type mice, as they were searching randomly in a first extinction test performed 2 hours after the probe trial, while wild type mice still showed a significant search compared to random (**Figure 1g**). This suggests that long-term memory was poorly consolidated in *Fus*^*ΔNLS/*+^ mice. Lastly, both genotypes were at random in a second extinction test (**Figure 1h**). Altogether, these data show that *Fus*^*ΔNLS/*+^ mice were able to learn, but displayed impaired long-term memory in agreement with a dysfunction in the cortical regions.

### Social disinhibition in *Fus*^ΔNLS/+^ mice

Marked changes in personality and social behavior, such as social withdrawal or social disinhibition, obsessive-compulsive behaviors, euphoria or apathy are common in subjects with behavioral variant (bv)FTD, a disease with pronounced FUS mislocalization^53-55^. Social deficits were also reported in progranulin haploinsufficient mice, a mouse model of FTD^56^. To determine whether *Fus*^*ΔNLS/*+^ mice have social behavioral deficits, we first performed the resident-intruder test specific for evaluating sociability in mice. Interestingly, 4 months old *Fus*^*ΔNLS/*+^ mice showed a trend towards longer interaction with the intruder mouse as compared with *Fus*^+*/*+^ mice (p= 0.07) (**Figure 2a**), that was significant at 10 months of age (**Figure 2b**) and persisted until 22 months of age (**Figure 2c**). To further characterize the social behavioral impairment, we used a modified version of the three-chamber social paradigm. After a first trial of habituation using an empty set up, a novel mouse is introduced in a side compartment. The interactions initiated by the test mouse with either the novel mouse or the empty cage was quantified. Of most relevance, across the three consecutive trials (Trial 2, 3 and 4), we observed that 10 months *Fus*^*ΔNLS/*+^ mice consistently interacted more with the novel mouse than *Fus*^+*/*+^ mice, further confirming social disinhibition (**Figure 2e**). This was not observed at 4 or 22 months of age **(Figure 2d-f).** Importantly, mice of both genotypes spent more time interacting with the novel mouse than with the empty cage, indicating that mice could recognize its conspecific. The interaction time gradually decreased in later trials, suggesting progressive loss of social interest in the novel mouse while it becomes a familiar mouse (**Figure 2d-f**). Similar findings of social disinhibition in both resident-intruder test and three chamber paradigms as a novel environment exclude the possibility that the observed increased social interactions resulted from locomotor hyperactivity in the home cage. Importantly, the olfactory function of *Fus*^*ΔNLS/*+^ mice was preserved, since results showed no differences between genotypes at 22 months of age in the time spent sniffing filter paper covered with either attractive scent (vanilla) or an aversive scent (2-methyl butyrate) (**Figure S2**). These findings together with absence of major motor phenotype at that age (**Figure S1a**,**b**) indicated that social behavior is specifically affected in *Fus*^*ΔNLS/*+^ mice. Taken altogether, behavioral analyses of *Fus*^*ΔNLS/*+^ mouse model unraveled locomotor hyperactivity, cognitive deficits and altered memory consolidation as well as selective impairment in sociability.

**Figure 2:**
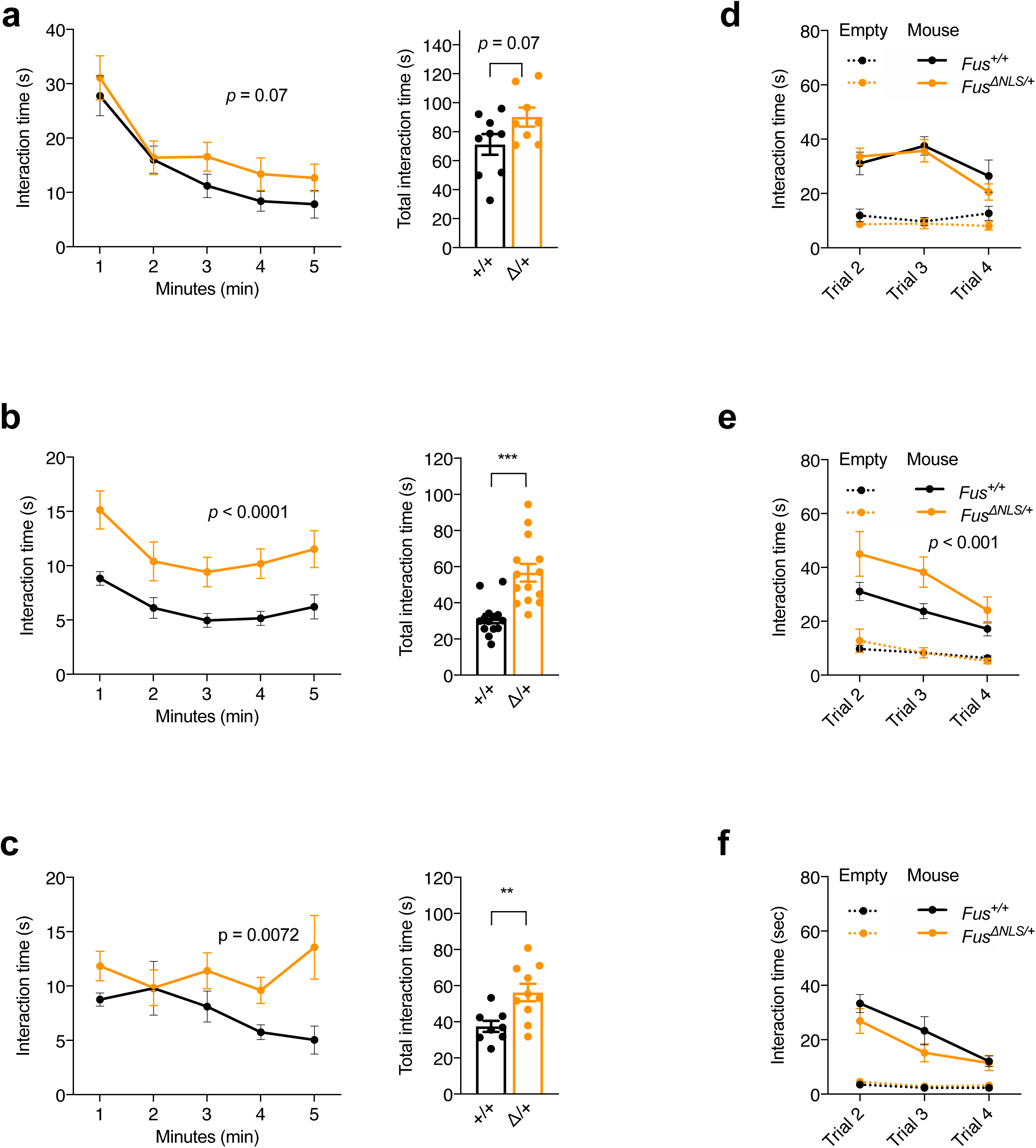
Social behaviour abnormalities in *Fus*^*ΔNLS*/+^ mice. **(a-c**) Line and bar graphs represent interaction time between resident (test) and intruder mice exclusively initiated by resident mouse per one minute intervals (line graphs, on the left) or per total time (bar graphs, on the right) during 5 minutes long resident intruder test in home cage for 4 (**a**), 10 (**b**) and 22 (**c**) months old *Fus*^+/+^ (black) and *Fus*^*ΔNLS*/+^ (orange) mice. Note that, young *Fus*^*ΔNLS*/+^ mice demonstrated a trend toward an increased social interest for intruder mouse (**a**) while older mice interacted with intruder significantly longer then *Fus*^+/+^ **(b**,**c**) showing age-dependent impairment of social behavior – disinhibition. All values are represented as mean +/- SEM. N=8-9 for 4 months, N=14 for 10 months and N=9-10 for 22 months. Two-way repeated measures ANOVA followed by Sidak *post hoc* test. p=0.07 (4 months), p<0.001 (10 months) and p=0.007 (22 months) for genotype.effect. Unpaired t-test for total time. p=0.07 (4 months), p<0.001 (10 months) and p=0.007 (22 months). (**d-f**) Line graphs represent sociability in three chamber test measured as interaction time with novel mice across three trials for *Fus*^+/+^(black) and *Fus*^*ΔNLS*/+^ (orange) mice at 4 (**d**), 10 (**e**) and 22 (**f**) months of age. Time exploring empty wired cage (object) across trials is represented as dashed lines. N=8-9 for 4 months, N=14 for 10 months and N=9 for 22 months. Three-way ANOVA with Newman Keuls *post-hoc* test for multiple comparisons. p=ns (4 months), p<0.001 (10 months) and p=ns (22 months) for genotype effect.

### Increased spontaneous neuronal activity in *Fus*^ΔNLS/+^ mice *in vivo*

As the observed behavioral changes are highly reminiscent of frontal lobe dysfunction, we next asked whether neuronal activity is altered within that brain area. Spontaneous neuronal activity was examined using *in vivo* two-photon calcium imaging (**Figure 3a-c**). Neurons in cortical layer II/III of the frontal cortex expressing the genetically encoded calcium indicator GCaMP6m were analyzed in 10 months old mice (**Figure 3b**). We observed a significant increase in spontaneous activity levels as seen in a larger fraction of active cells in *Fus*^*ΔNLS/*+^ (**Figure 3d**, p=0.013, student’s t-test, n=14 experiments in *Fus*^*ΔNLS/*+^ mice and n=10 experiments in *Fus*^+*/*+^ mice). Moreover, we observed a right shift in the distribution of the transient amplitudes (**Figure 3e**, p<10^−16^, KS test) and in the transient frequency (**Figure 3f**, p<10^−8^, KS test) in *Fus*^*ΔNLS/*+^ mice compared to their *Fus*^+*/*+^ littermates (n=631 neurons in 5 *Fus*^+/+^ mice; n=855 neurons in 6 *Fus*^ΔNLS/+^ mice). Taken together, these results demonstrate a strong increase in neuronal activity *in vivo* within the upper layers of frontal cortex of *Fus*^ΔNLS/+^ mice.

**Figure 3.**
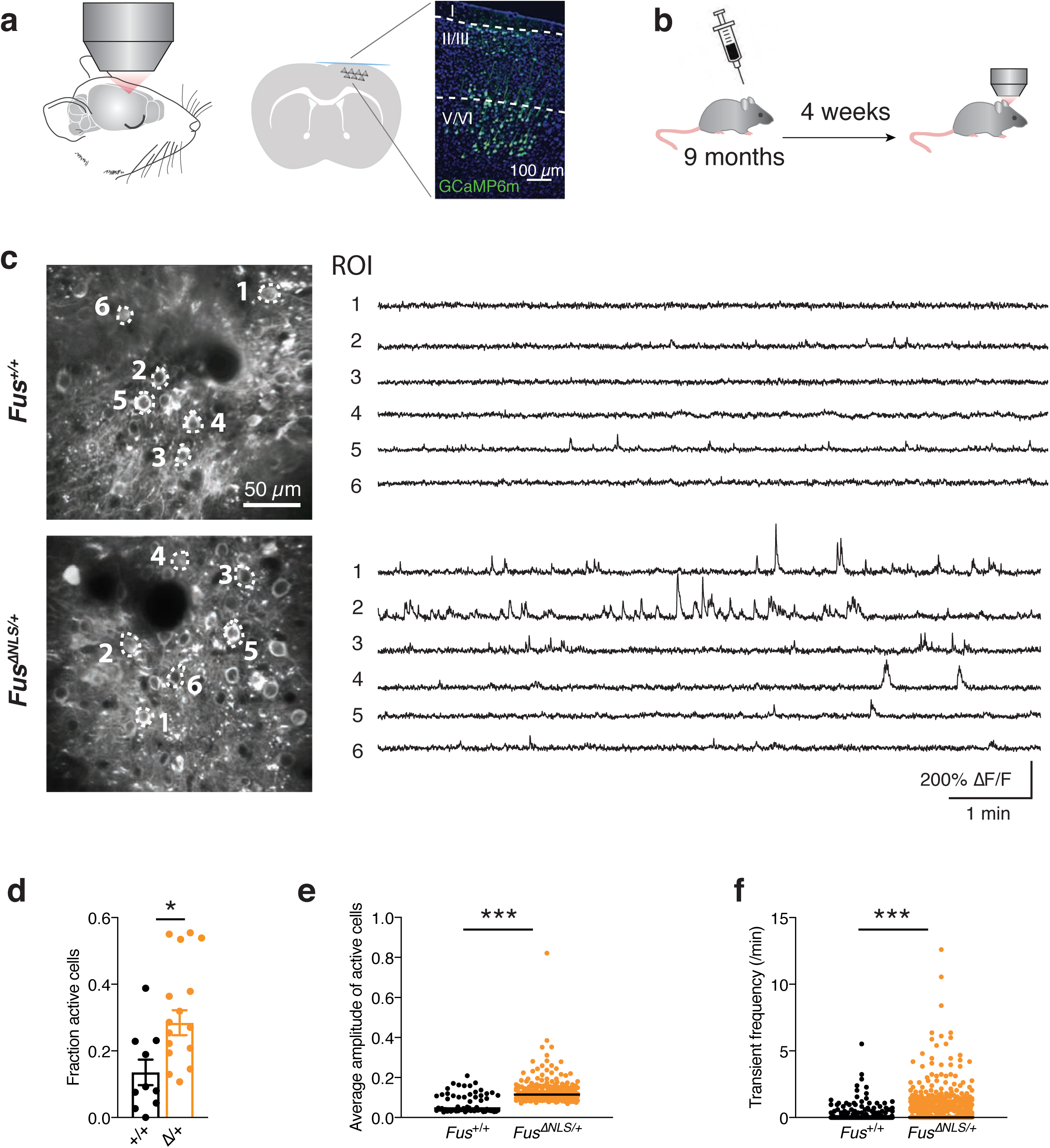
Assessment of neuronal activity in *Fus*^*ΔNLS/*+^ mice *in vivo*. **(a)** Neuronal activity is monitored in frontal cortex in anesthetized mice. Scheme of coronal section, indicating expression of GCaMP6s in cortex assessed through a cranial window. Magnified view of imaged cortical area demonstrates neuronal expression of GCaMP (green) in all cortical layers. **(b)** Time line of experiments. Mice are injected with AAV2/1-hsyn-GCaMP6m into frontal cortex and implanted with a cranial window at 9months of age. In vivo imaging commences 4 weeks after implantation. **(c)** Representative examples (average projections) of field of views (FOV) imaged in both mouse lines are shown together with fluorescent calcium traces of selected regions of interest (ROIs). **(d-f)** The fraction of active cells per FOV (**d)** as well as **(e)** the frequency and **(f)** the average amplitude of calcium transients of each ROI is increased in *Fus*^*ΔNLS*/+^ mice. Data are individual FOVs (d; 14 FOVs in 6 *Fus*^*ΔNLS*/+^ and 10 FOVs in 5 *Fus*^+/+^ mice) or individual ROIs (e-f; 855 ROIs in *Fus*^*ΔNLS*/+^ and 631 ROIs in *Fus*^+/+^) superimposed by the mean +/- SEM (d) or the median e, f). * p < 0.05, *** p < 0.001

### *Fus*^ΔNLS/+^ mice show ventricle enlargement but lack obvious neuronal loss

We then sought to understand the etiology of behavioral and electrophysiological abnormalities in *Fus*^*ΔNLS/*+^ mice. Using MRI, we observed an increased ventricle volume in 12 months old *Fus*^*ΔNLS/*+^ mice, but roughly preserved cortex volume (**Figure 4a-e**). In order to determine whether the observed phenotypes could be linked to neurodegeneration in the cortex, we performed brain histology in *Fus*^*ΔNLS/*+^ mice at both 10 and 22 months of age. Cortical cytoarchitecture appeared preserved in *Fus*^*ΔNLS/*+^ mice, with normal lamination and no cortical thinning. The density of NeuN positive neurons in the frontal cortex was similar between *Fus*^*ΔNLS/*+^ mice and their wild type littermates at 10 and 22 months of age (**Figure 4f-g**). These data suggest that the behavioral phenotypes observed in *Fus*^*ΔNLS/*+^ mice result rather from neuronal dysfunction than from generalized neuronal loss in the cortex.

**Figure 4.**
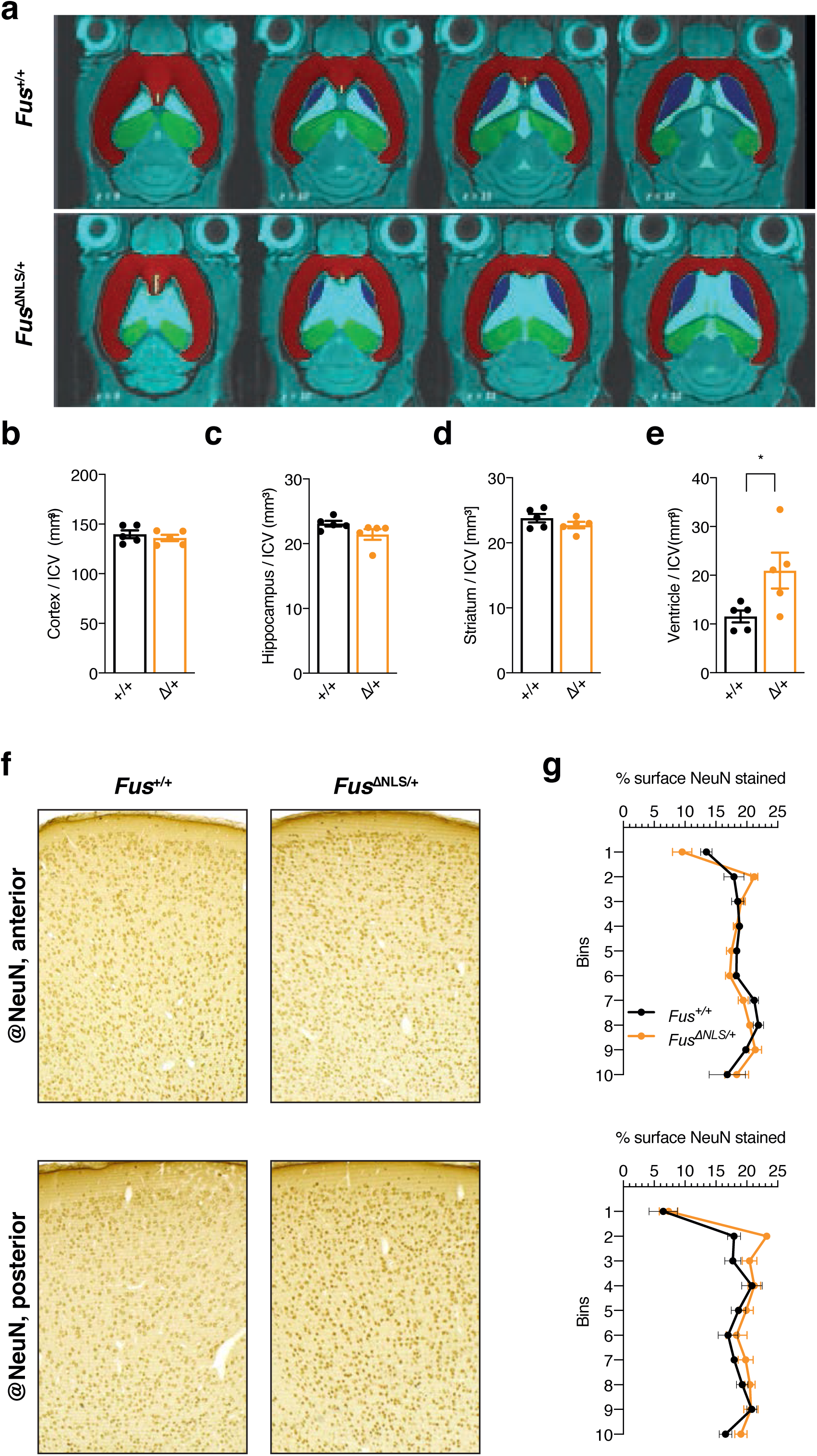
Structural and histological brain analysis of *Fus*^+/+^ and *Fus*^*ΔNLS*/+^ mice. **(a)** Volume determination by Tissue Classification Software (TCS) in 12 months old *Fus*^*ΔNLS/*+^ (**lower row**) and *Fus*^+*/*+^ (**upper row**). Striatum (blue), cortex (red), hippocampus (green), and ventricles (light blue) were identified in 30 slices. **(b-e)** Comparison at the group level of *Fus*^*ΔNLS/*+^ vs *Fus*^+*/*+^, normalized to ICV; significance level was p-value < 0.05 *. ICV = intracranial volume. **(f)** Representative image of NeuN immunohistochemistry at 22 months of age in *Fus*^+/+^ or *Fus*^ΔNLS/+^ mice, in anterior and posterior regions of the M1/M2 cerebral cortex **(g)** Distribution of NeuN+ neurons in *Fus*^+/+^ (black) or *Fus*^ΔNLS/+^ (orange) mice, in anterior and posterior regions of the M1/M2 cerebral cortex.

### Transcriptome of *Fus*^ΔNLS/+^ cortex points to defects in inhibitory transmission

To understand the molecular basis of neuronal dysfunction, we then performed RNAseq on frontal cortex of 5 and 22 months old *Fus*^*ΔNLS/*+^ mice and their wild type littermates. Principal component analysis showed a clear separation between *Fus*^*ΔNLS/*+^ mice and their wild type littermates at 22 months of age, but clustering was imperfect at 5 months of age suggesting an exacerbation of the transcriptional differences between genotypes with age (**Figure 5a**). Statistical comparison identified that most altered genes were increased in expression at 5 months of age (644 genes significantly upregulated vs 289 downregulated at 5 months of age, FDR < 0.1, Log2 fold change > 0.5 and < -0.5), while the opposite was observed at 22 months of age (81 genes upregulated and 398 genes downregulated, **Figure 5b**). When we analyzed all differentially expressed genes using FDR < 0.1, 182 genes were commonly regulated at 5 and 22 months of age, which were selectively enriched in genes functionally involved in GABA synthesis, GABAergic synapse and neuropeptide according to gene ontology analysis (**Figure 5c, Supplementary Table 1**). Interestingly, GABA-related genes tended to be upregulated at 5 months of age and downregulated at 22 months of age (**Figure 5d-e, Supplementary Table 2**). To determine whether the expression of FUS was intrinsically required for GABAergic neurons, we compared the transcriptome at both ages with previous results obtained in embryonic *Fus*^*-/-*^ brain^10^. As shown in the Chord diagram of **Figure 5f**, the 3 enriched GO terms (GABAergic synapse, neuronal differentiation and neuronal system) were consistently enriched across the different mouse models and developmental stages/age cohorts, suggesting a tight relationship between FUS function and GABA (**Figure 5f**). Of note, this enrichment was not observed in embryonic *Fus*^*ΔNLS/*Δ*NLS*^ brains^10^ (data not shown), suggesting that the effect of the ΔNLS mutation occurs later than the complete ablation of the *Fus* gene. Thus, expression of multiple genes related to inhibitory neurotransmission was altered in the frontal cortex of *Fus*^*ΔNLS/*+^ mice.

**Figure 5:**
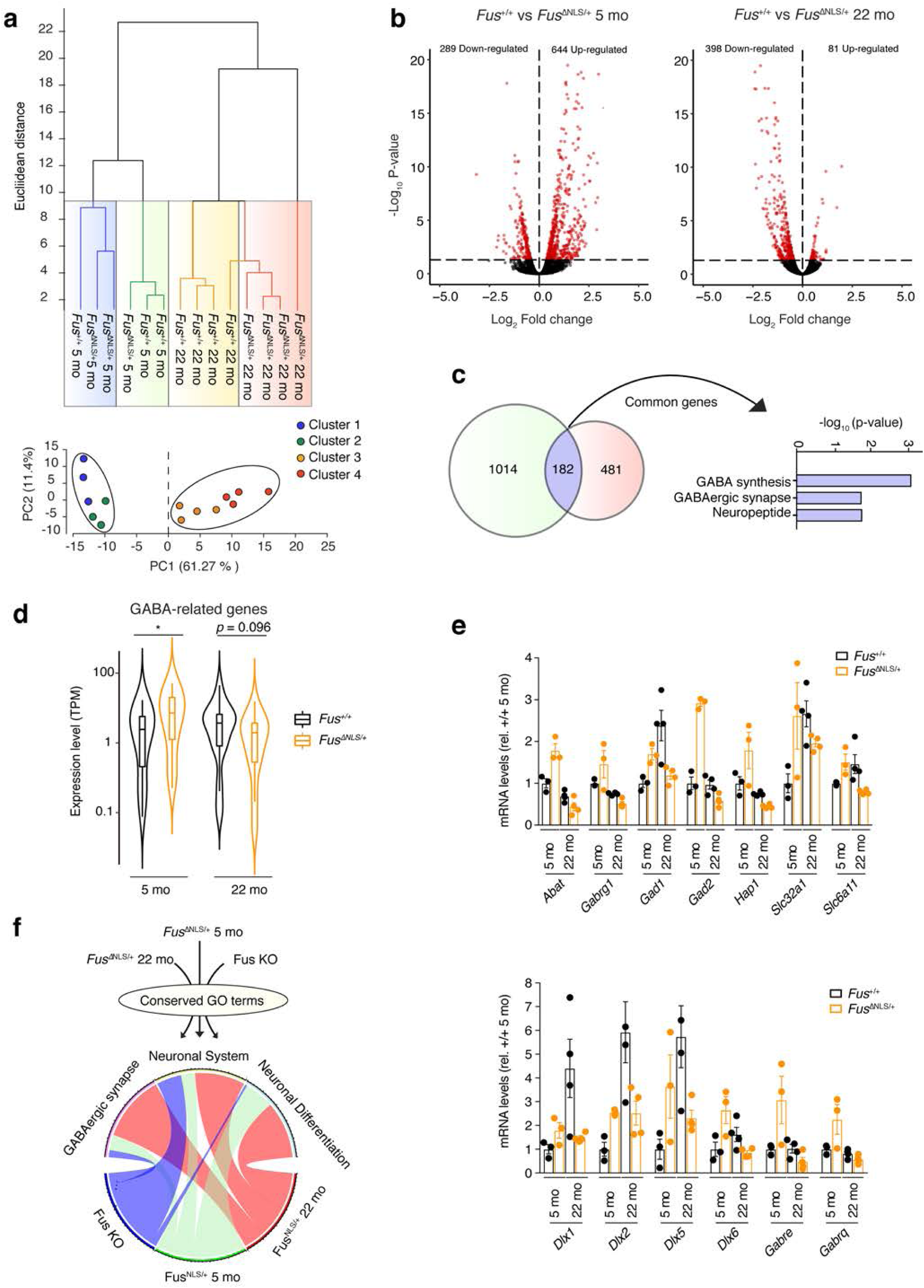
Longitudinal transcriptome analysis points to defects in inhibitory synapses in *Fus*^*ΔNLS*/+^ mice. **(a)** Hierarchical clustering and principal components analysis (PCA) shows that the first PCA differentiates between 5 months and 22 months old mice while PC2 distinguishes between *Fus*^+/+^ and *Fus*^ΔNLS/+^ mice only in young animals. **(b)** The volcano plot shows a clear asymmetric profile in young *Fus*^ΔNLS/+^ animals with more genes being up-regulated (644 genes) than down-regulated (244 genes). Conversely, in old *Fus*^ΔNLS/+^ mice, we observed that more genes are down-regulated (398 genes) than up-regulated (81 genes) after multiple correction (FDR < 0.1, Log2 fold change > 0.5 and < -0.5) **(c)** We identified 182 overlapping genes between 22 months and 5 months old mice when we analyzed all differentially expressed genes using FDR < 0.1 as criterion. GO enrichment analysis of this overlapping gene set identifies GABA-related terms associated with the GABAergic synapse (Hypergeometric test p < 0.01) **(d)** Histogram of TPM values of the GABA-related genes identified in (c) shows an up-regulation in 5 months old *Fus*^ΔNLS/+^ mice compared to their wild type littermates (Welch corrected T-test : t = 2.123, df = 28.64, p = 0.048). Conversely, the same set of genes show a trend towards a down-regulation in 22 months old *Fus*^ΔNLS/+^ mice compared to their wild-type littermates (Welch corrected T-test : t = 1.73, df = 23.64, p = 0.096) **(e)** Histograms with individual GABA-related genes in the adult prefrontal cortex. We observe that the GABA-related genes are up-regulated in 5 months old *Fus*^ΔNLS/+^ mice while they are downregulated in 22 months old *Fus*^ΔNLS/+^ mice compared to their WT litternates. Statistics are presented in Supplementary Table 3. **(f)** Significant GO terms (FDR < 0.01) were merged among three independent models carrying a loss or gain of function mutation. Conserved GO terms were then plotted using a chord diagram (custom Rscript). We observed that the most conserved significant GO terms in 5 months, 22 months old and Fus KO mice are related to the GABAergic system highlighting its role in FUS-associated pathology.

### Defects in inhibitory neurons in *Fus*^*ΔNLS/*+^ mice

In order to study a potential defect in inhibitory neurons in *Fus*^*ΔNLS/*+^ cortex, we focused on parvalbumin positive (PV) interneurons. Indeed, in addition of being the largest group of inhibitory interneurons in the cortex, evidence suggests that the transcriptional changes observed could be mainly driven by altered PV neurons. First, the *Pvalb* gene, encoding parvalbumin, was downregulated at 5 months of age and upregulated at 22 months of age in *Fus*^*ΔNLS/*+^ cortex (**Figure S3)**. Second, *Dlx5* and *Dlx6*, encoding two critical transcription factors involved in PV neuronal development or *Gabrg1*, a GABA receptor subunit mostly expressed in a subset of PV neurons^57,58^, were among the 182 genes commonly regulated at both ages (**Figure 5d**). Last, markers of other inhibitory neuron populations, such as *Sst, Htr3a, Npy, Cck* or *Vip*, were either unchanged at both ages or downregulated only at early time points (**Figure S3**). Using immunohistochemistry, we did not detect differences in the number or layering of PV neurons in the frontal cortex of *Fus*^*ΔNLS/*+^ mice either at 10 or 22 months of age (**Figure S4a-c**). To determine whether FUS was mislocalized in PV neurons, we performed double immunostaining for FUS and parvalbumin, and determined the nuclear/cytoplasmic ratio selectively in PV neurons. As shown in **Figure S4d-e**, cytoplasmic FUS staining was increased in *Fus*^*ΔNLS/*+^ as compared with *Fus*^+*/*+^ PV positive neurons.

Intriguingly, FUS cytoplasmic staining increased with age in wild type PV interneurons, but remained significantly lower than in *Fus*^*ΔNLS/*+^ neurons. Quantitative ultrastructural analysis of (symmetric) inhibitory synapses in layers II/III of the frontal cortex demonstrated that *Fus*^*ΔNLS/*+^ inhibitory boutons were larger in size (**Figure 6a-c**), with longer active zone (**Figure 6d**), increased vesicle numbers (**Figure 6e**) and that the mean distance of vesicles to the active zone was longer than in wild type synapses (**Figure 6f**). To further explore morphological changes occurring at inhibitory synapses, we quantified the density and the cluster size of three inhibitory synaptic markers: the GABA transporter VGAT localized at the presynaptic site^59^ and two receptors specifically expressed at the postsynaptic site of GABA monoaminergic synapses^60^, the postsynaptic scaffold protein Gephyrin^61^ and the GABA_A_ receptor containing α3 subunit (GABA_A_Rα3). Pictures were acquired in the layer 1 of the cortex to allow imaging of inhibitory synapses located on the apical dendrites of pyramidal neurons^62^. Consistent with the observed ultrastructural abnormalities, a significant decrease in all markers for inhibitory synapses was identified (**Figure 6g-i**) (*Fus*^+*/*+^ vs *Fus*^*ΔNLS/*+^, VGAT, p=0.0464; GABA _A_ Rα3, p=0.0217; Gephyrin, p=0.0043). This decrease in density was associated with a decrease in the size of the clusters for VGAT, GABA _A_ Rα3 and Gephyrin (*Fus*^+*/*+^ vs *Fus*^*ΔNLS/*+^, for VGAT, GABA _A_ Rα3 and Gephyrin, p<0.0001), suggesting a functional impairment of the remaining synapses. Altogether, these results demonstrate that cortical inhibitory interneurons are affected in *Fus*^*ΔNLS/*+^ mice, which could underlie the observed neuronal hyperexcitability (**Figure 3**).

**Figure 6:**
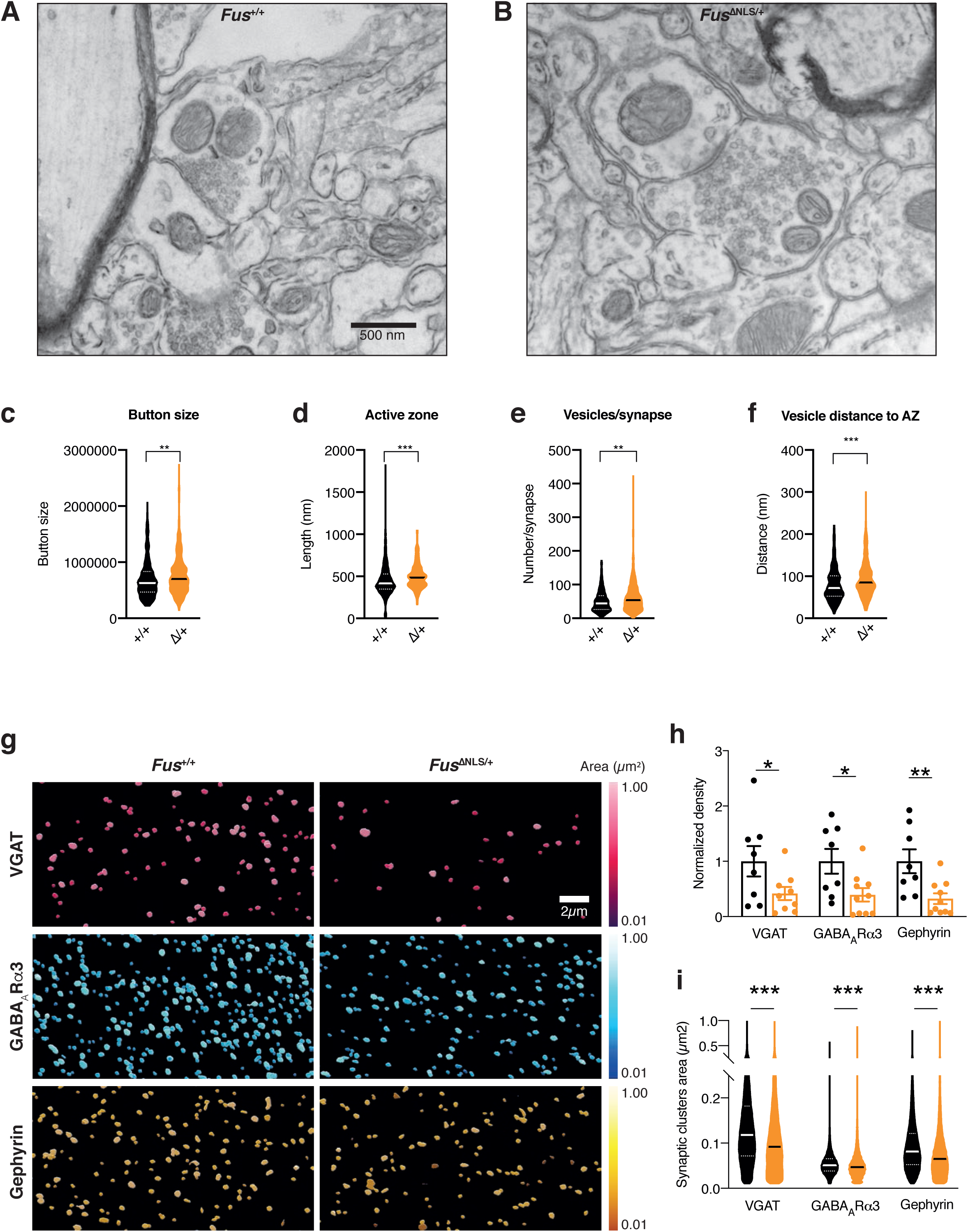
defects in inhibitory synapses in 22 months old *Fus*^*ΔNLS*/+^ mice. **(a-b)** Representative image of transmission electron microscopy in *Fus*^+/+^ (a) or *Fus*^ΔNLS/+^ (b) layer II/III of the motor cortex at 22 months of age. Inhibitory synapses were identified as containing ≥1 mitochondria on each side of the synapse. **(c-f)** Violin plot showing the distribution of bouton size (c), the length of active zone (d), the number of vesicles per synapse (e) and the distance of individual vesicles to the active zone (f) in inhibitory synapses of *Fus*^+/+^ (black) or *Fus*^ΔNLS/+^ (orange) mice. **(g)** Representative images of GABAARα3, Gephyrin and VGAT intensity at 22 months of age coded by area size (Imaris) **(h)** Bar graphs representing the density analysis for VGAT, GABAARα3 and Gephyrin comparing *Fus*^+/+^ vs *Fus*^ΔNLS/+^ mice. (*Fus*^+/+^ vs *Fus*^ΔNLS/+^, Mann-Whitney test, VGAT, p=0.0464; GABAARα3, p=0.0217; Gephyrin, p=0.0043) **(i)** Violin plot representing the analysis of the clusters size for VGAT, GABAARα3 and Gephyrin comparing *Fus*^+/+^ vs *Fus*^ΔNLS/+^ mice. (*Fus*^+/+^ vs *Fus*^ΔNLS/+^, Unpaired t-test, for VGAT, GABAARα3 and Gephyrin, p<0.0001)

### Abnormal synaptosomal RNAs in *Fus*^*ΔNLS/*+^ cortex

To determine whether the observed phenotypes could be linked to a disrupted function of FUS at the synapse, we performed synaptosomal fractionation of the frontal cortex from 5 months old *Fus*^*ΔNLS/*+^ mice. Obtained fractions were enriched in the synaptophysin protein (**Figure 7a-b**, and **Figure S5** for uncropped western blots) and depleted in the nuclear lncRNA *Malat* (**Figure 7c**), consistent with synaptic enrichment. In these fractions, we observed strongly enriched levels of the FUS protein as detected using N-terminal antibodies, but not using NLS targeting antibodies (**Figure 7a-b**), suggesting that mutant FUS protein preferentially accumulated in synaptosomal fractions as compared to the wild type protein. FUS is known to bind a number of mRNAs, including *Fus* mRNA itself, as well as mRNAs important for (inhibitory) synaptic function such as *Nrxn1* or *Gabra1*^*34*^. Consistently, we observed increased levels of these 3 mRNAs in synaptosomal fractions of *Fus*^*ΔNLS/*+^ mice (**Figure 7c**). This enrichment was relatively selective as mRNAs encoding other GABA related proteins, such as *Gphn, Gabra3, Vgat, Gabrb1, Gabrg1* and *Gad2* or acetylcholine related proteins such as *Chrn2b*, were not enriched in the synaptosomal fraction by the *Fus* mutation. We also observed a slight increase in *Chrna7* mRNA in *Fus*^*ΔNLS/*+^ synaptosomes. Collectively, our data point to defects in inhibitory synapses, that might cause the increased spontaneous neuronal activity and subsequent widespread behavioral abnormalities in *Fus*^*ΔNLS/*+^ mice.

**Figure 7:**
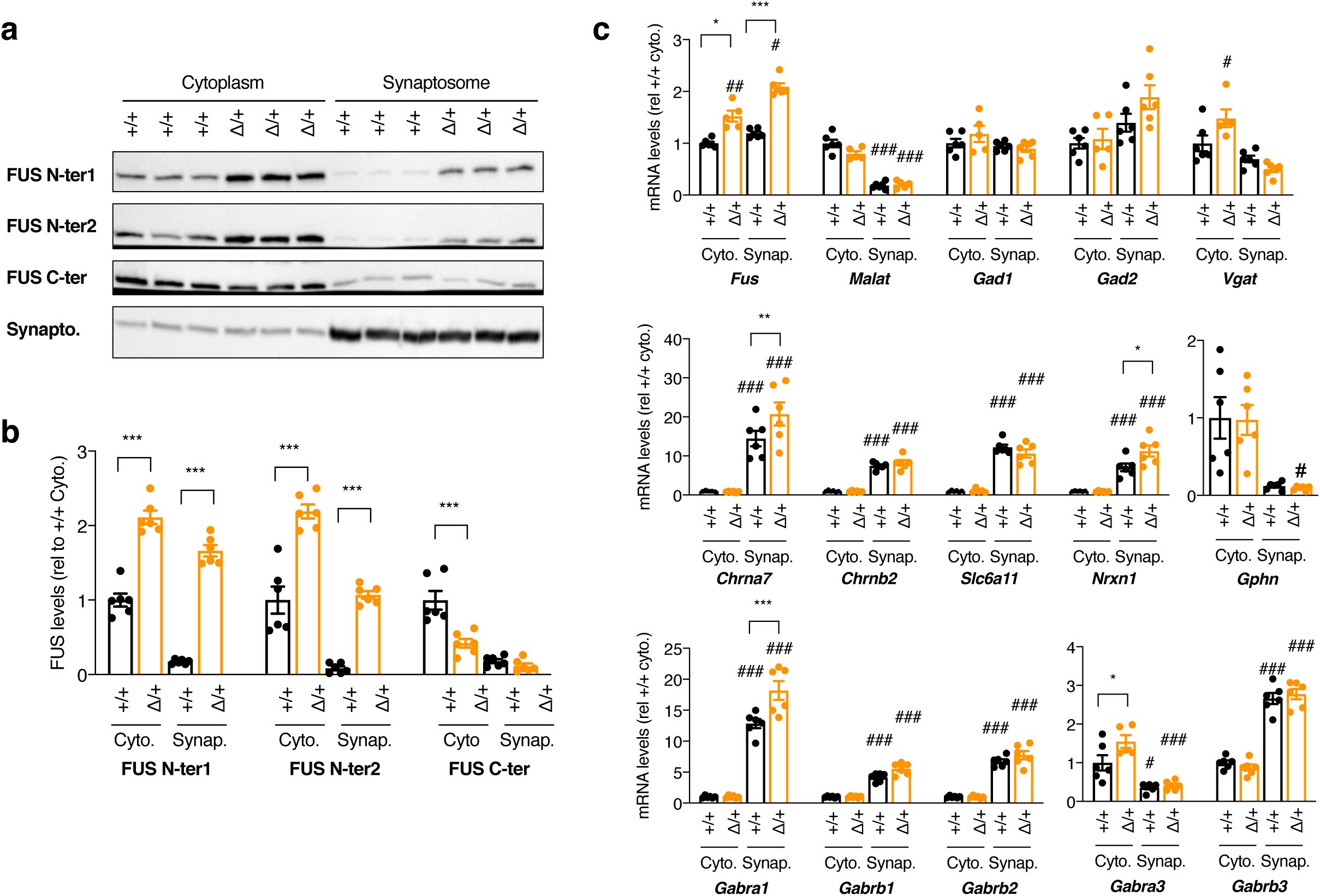
FUS accumulates in synaptosomes of *Fus*^*ΔNLS*/+^ mice and alters synaptosomal levels of a subset of its targets. (**a-b**) Representative western blot images (**a**) and respective quantifications (**b**) of cytoplasmic (left) or synaptosome (right) extracts from *Fus*^+/+^ (+/+) or *Fus*^ΔNLS/+^ (Δ/+) mice (4 months of age, 3 individual mice presented, 6 mice in total analyzed per genotype) using two antibodies recognizing the N-terminal part of the FUS protein (FUS N-ter1 and FUS N-ter2), the C-terminal part of FUS (encoding the NLS, FUS C-ter) or synaptophysin protein to show enrichment in synaptic proteins in the synaptosome fraction. Please note that the 3 FUS western blots were run on independent gels, to avoid stripping and reprobing on the same membrane for the same protein. Each of these gels were controlled for equal loading using StainFree markers, that are provided in Figure S5 with uncropped western blot images. (c) mRNA levels of the indicated genes in RNAs extracted from cytoplasmic (Cyto.) or synaptosome (Synap.) extracts from *Fus*^+/+^ (+/+) or *Fus*^ΔNLS/+^ (Δ/+) frontal cortex from 4 months old mice as assessed using RT-qPCR. All quantifications are presented relative to the +/+ cytoplasmic RNA levels set to 1. *, p<0.05 vs corresponding wild type fraction; ***, p<0.001 vs corresponding wild type fraction; #, p<0.05 vs corresponding cytoplasmic fraction of the same genotype; ###, p<0.001 vs corresponding cytoplasmic fraction of the same genotype; Two-way repeated measures ANOVA with Tukey *post hoc* test.

## Discussion

In this study, we show that knock-in mice with cytoplasmic accumulation of FUS display widespread behavioral alterations, beyond motor symptoms. We further determine that FUS mislocalization leads to increased spontaneous neuronal activity in the cortex, indicative of neuronal hyperexcitability, that is associated with alteration in inhibitory synapses. Last, we show that FUS mutation alters FUS synaptic content and modifies synaptic levels of a subset of its RNA targets, possibly underlying the observed phenotypes.

The recognition that FUS mislocalization is a widespread pathological event in sporadic ALS, but also in many other neurological diseases prompted us to investigate the behavioral phenotype of *Fus*^ΔNLS/+^ mice. While motor defects can be detected as early as 6 months of age and motor neuron degeneration is not detected before 18-22 months of age, we observed an early and sustained spontaneous locomotor hyperactivity in *Fus*^ΔNLS/+^ mice. This hyperactivity did not appear to be associated with increased anxiety as open field behavior or performances in the light/dark box test were normal. In addition, we observed various defects in executive functions, including impaired remote long-term memory, and abnormal social interactions. Hyperactivity and social and executive dysfunctions have been previously documented in other mouse models of ALS/FTD. It is noteworthy for instance that transgenic overexpression of mutant FUS leads to hyperactivity and cognitive deficits^63^. Similar abnormalities are also observed in TDP-43 knock-in mice^64^, C9ORF72 BAC transgenic mice^65^ or Chmp2b transgenic mice^66^, suggesting that ALS mutations commonly lead to various behavioral alterations in mouse models, that are dominant over motor dysfunction. The observation of such phenotypes in mouse models is relevant to the existence of widespread cognitive and executive dysfunction in ALS patients^67,68^ and support the clinical overlap between ALS and FTD^69^.

The deficits in executive functions and social behaviors that we observe in *Fus*^ΔNLS/+^ mice are particularly relevant for FTD. Pathology of FUS and the other FET proteins (TAF15 and EWRS1) is a hallmark of a subset of FTD cases (FTD-FET cases), and these patients also display prominent atrophy of the caudate putamen^24-26^, in line with our finding of an increased volume of the third ventricle observed by MRI in *Fus*^ΔNLS/+^ mice. In FTD-FET cases, FUS pathology is associated with nuclear clearance of the FUS protein in neurons with FUS aggregates, although this nuclear clearance is not as pronounced as in cases with TDP-43 pathology^40^. Importantly, the FUS protein is accompanied by several other proteins in FTD-FET pathological aggregates, including TAF15 and EWSR1, two other FET proteins, as well as Transportin 1^12,27-30^. Thus, the disease in FTD-FET patients could be driven by several non-mutually exclusive mechanisms, including cytoplasmic accumulation and/or aggregation of FUS, nuclear clearance of FUS and/or aggregation of co-deposited pathological proteins. Previous studies indicated that complete loss of FUS could be sufficient to lead to FTD like symptoms in mice, and this was consistent with the role of FUS in controlling the splicing of mRNAs relevant to FTD, such as *MAPT*, encoding the TAU protein, or in the stability of mRNAs encoding relevant synaptic proteins such as GluA1 and SynGAP1^35-39^. In *Fus*^ΔNLS/+^ mice, there is, however, a limited loss of nuclear FUS immunoreactivity^10,41^ and no obvious FUS aggregates, ruling out that these pathological events might play a major role in the observed behavioral alterations. The quasi-normal levels of FUS in the nucleus are explained by the existence of potent autoregulatory mechanisms, able to largely buffer the effect of the mutation on nuclear FUS levels. Mislocalization of either TAF15 or EWSR1 is also unable to account for behavioral abnormalities as both of these proteins show normal localization in *Fus*^ΔNLS/+^ neurons, as well as ALS-FUS patients^30^. Together, our results show that FUS mislocalization alone is sufficient to trigger behavioral symptoms, and suggest that this might be a major driver of disease pathophysiology in FTD-FET patients. Importantly, this does not exclude that at later stages of disease progression, loss of nuclear FUS function might occur as a result of saturation of autoregulatory mechanisms, thereby exacerbating neurological symptoms.

A major finding of this study is that *Fus*^ΔNLS/+^ mice develop morphological and ultrastructural synaptic defects. The combination of locomotor hyperactivity with social deficits, as observed in *Fus*^ΔNLS/+^ mice, is commonly observed in various mouse models with synaptic defects. For instance, mouse models of autism spectrum disorders, such as mice ablated from the ProSAP/Shank proteins^70,71^, display similar behavioral alterations. Our results point to a major defect in inhibitory synapses. This conclusion is supported by at least three main results: First, transcriptome of the cerebral cortex shows that genes related to GABAergic synapses are affected, being coordinately upregulated in young adults, and downregulated at later stages. Second, inhibitory synapses are ultrastructurally abnormal, with increased size, increased number of vesicles and increased distance between vesicles and the active zone. Third, the density of inhibitory synapses as well as the clusters size of three typical markers of inhibitory synapses (VGAT, GABA_A_Ra3 and Gephyrin) are decreased. Our data suggest that both the pre- and post-synaptic part of inhibitory synapses are affected by the *Fus* mutation. Indeed, the decrease in the density of Gephyrin likely reflects a disorganization of the postsynapse density^72^, that could be caused by decreased GABAR activity^73-75^. Decreased VGAT density, as well as increased bouton size or vesicle disorganization further suggest impairment of presynaptic GABAergic terminals. On its own, decreased VGAT density might explain a reduction of synapses throughout the cortical layers^76^ and lead to impaired loading of GABA in the presynaptic vesicles^59^. Importantly, Sahadevan, Hembach and collaborators performed studies in *Fus*^ΔNLS/+^ mice at earlier ages and also observed early defects of inhibitory synapses, as early as 1 month of age and progressive at 6 months of age (Sahadevan, Hembach et al, co-submitted manuscript). It is interesting to note that the disruption of inhibitory synapses is able to explain most of the observed behavioral and electrophysiological phenotypes observed in *Fus*^ΔNLS/+^ mice. Illustrating this, loss of *Gabra1*^77^, or of *Gabra3*^78^ are sufficient to lead to locomotor hyperactivity and the FUS target *Nrxn1*, encoding a critical actor in the formation of GABAergic and glutamatergic synapses^79^, is critical in locomotor activity and social behavior in mice^80,81^. Indeed, deletion of all three neurexins from PV neurons is causing a decrease in the number of synapses in this neuronal type^82^, in a manner similar to what is observed in *Fus*^ΔNLS/+^ mice.

Is a specific subpopulation of inhibitory neurons selectively affected in *Fus*^ΔNLS/+^ mice? The transcriptome analysis showed that markers of PV interneurons, such as *Pvalb*, were more severely affected than markers of other interneurons (eg *Sst)*. Furthermore, several genes critically involved in PV neuronal development were also affected in *Fus*^ΔNLS/+^ mice. The involvement of PV neurons was also consistent with previous studies in TDP-43 knock-in mice, showing PV interneurons impairment^64^, and in TDP-43 transgenic mice displaying degeneration of hippocampal PV positive interneurons^83^. Functional impairment of PV interneurons might represent a unifying theme in ALS pathophysiology, as multiple electrophysiological studies demonstrate hypoexcitability of PV neurons in SOD1 and TDP-43 transgenic mouse models of ALS^84-86^. Others, however, found PV interneurons to be unaltered presymptomatically and to turn hyperexcitable during the symptomatic phase in the same SOD1^G93A^ mouse model^87^. In either case, those changes in PV excitability were always accompanied by hyperexcitability of layer V pyramidal neurons^84-87^. These findings in mouse models nicely recapitulate human ALS pathology, in which cortical hyperexcitability is a frequent and, most importantly, early finding in familial and sporadic cases, including FUS mutation carriers^16,88^. In line with these findings, we also observed a pronounced increase in spontaneous neuronal activity *in vivo*, which is highly indicative of hyperexcitable pyramidal neurons. While we cannot rule out cell autonomous alterations affecting the intrinsic excitability of pyramidal neurons, our histological, ultrastructural and transcriptomic data strongly argue for defective inhibitory neurotransmission by PV interneurons. In summary, our results, along with others, support that dysfunction of cortical PV interneurons contribute to defects in ALS and FTD. Importantly, while we observe molecular and structural defects in inhibitory neurons, we did not observe loss of PV cell bodies in *Fus*^ΔNLS/+^ mice, suggesting that the major defects are synaptopathies rather than neuronal loss, consistent with other studies^51^. Altogether, our results identify a role for FUS in regulating GABAergic synapse structure and function. Since other major classes of inhibitory interneurons^57^ were not investigated, we cannot exclude that SST neurons or HTR3A neurons are also affected, although to a lesser extent than PV neurons. Furthermore, our work does not exclude that FUS regulates also glutamatergic synapses. On the contrary, Sahadevan, Hembach et al. (co-submitted manuscript) provide evidence that FUS is also critically involved in glutamatergic synaptogenesis, at least during development, as did previous studies^89^. Further work is required to disentangle the causes and consequences of GABAergic and glutamatergic impairment, and their respective mechanisms.

How can mutant FUS regulate inhibitory synaptic structure? We observe that the loss of the FUS NLS leads to an increased level of the mutant protein in purified synaptosomes. These results are entirely consistent with results from Sahadevan, Hembach et al. These authors identified a number of FUS synaptic RNA targets, and a subset of these were also increased in synaptosomes of *Fus*^ΔNLS/+^ mice. Interestingly, in both studies, a number of FUS synaptic targets are not modified in *Fus*^ΔNLS/+^ synaptosomes, including some related to GABAergic neurons. Besides mRNA trafficking, FUS accumulation in synaptosomes might alter local synaptic translation^44,90^, thereby modifying protein levels. This is in particular the case for GABRA3, whose mRNA, a strong FUS synaptic target (Sahadevan, Hembach et al. co-submitted manuscript), is not increased in *Fus*^ΔNLS/+^ synaptosomes, while GABRA3 protein is decreasing with age at synaptic sites (this manuscript and Sahadevan, Hembach et al. co-submitted manuscript). Further work should focus on determining whether FUS might also regulate synaptic translation of specific proteins involved in inhibitory transmission.

In summary, we show here that cytoplasmic accumulation of FUS leads to a major synaptopathy in inhibitory neurons, that is accompanied by consistent behavioral and electrophysiological phenotypes. The identification of the mechanisms downstream of FUS synaptic action might lead to efficient therapeutic strategies for FUS related neurodegenerative diseases.

## Materials and methods

### Mouse models and behavioral analyses

Wild type (*Fus*^+*/*+^) and heterozygous (*Fus*^ΔNLS/+^) mice on a pure genetic background (C57/Bl6), generated as described previously^10^, were bred and housed in the central animal facility of the Faculty of medicine of Strasbourg, with a regular 12-h light and dark cycle (light on at 7:00 am) under constant conditions (21 ± 1 °C; 60% humidity). Standard laboratory rodent food and water were available *ad libitum* throughout all experiments. Mice were genotyped by PCR of genomic DNA from tail biopsies as described previously^10^. Mouse experiments were approved by local ethical committee from Strasbourg University (CREMEAS) under reference number AL/27/34/02/13 (behavior), by the Government of upper Bavaria (license number Az 55.2-1-54-2532-11-2016, two photon microscopy) and by "Regierungspräsidium Tübingen” (animal license number 1431, MRI). Behavioral tests were done during the light phase (between 9 am and 5 pm) of their light/dark cycle except for indicated experiments. Male mice of 4, 10 and 22 months of age were subjected to behavioral studies and data were analyzed blind to genotypes.

### Spontaneous locomotor activity in the home cage – actimetry

Home cage activity was assessed as described previously^91^. Briefly, the mice were placed individually in large transparent Makrolon cages (42 × 26 × 15 cm) adapted to the shelves of the testing device (eight cages/shelve). Two infrared light beams, passing through each cage, were targeted on two photocells, 2.5 cm above the cage floor level and 28 cm apart. The number of cage crossing was recorded automatically and was used to determine or score the spontaneous locomotor activity. The experiment began at 17.00 pm and after 2 hours of habituation continued for 3 consecutive days for a complete 24 h nictemeral cycle (12h dark and 12h light).

### Open field

The general exploratory locomotion and anxiety in a novel environment were tested during 15 min long sessions in the open field arena (72 × 72 × 36 cm) located in a test room and lit by a 600 lux for background lighting, according to published protocol^92^. The open maze was divided by lines into sixteen squares (18 × 18 cm). Each mouse was placed in the center of the arena and allowed to freely move while being video recorded. The recorded data were analyzed offline with EthoVision XT software system (Noldus Information Technology). The time spent in the center (four central quadrants) vs. the perimeter (12 peripheral quadrants) was used to measure anxiety, while the total distance traversed in the arena and average moving speed (mean velocity) was used to evaluate locomotor activity. For each mouse a movement heat map and trajectory tracking map that are representing a corresponding locomotor activity were made independently.

### Dark/light box test

The light/dark box apparatus consisted of two Poly-Vinyl-Chloride (PVC) compartments of equal size (18.5 × 18.5 × 15 cm) one opaque and the other transparent, connected through an opaque tunnel (5 × 5.5 × 5 cm). The illumination of the transparent compartment was set at 400 lux. Each mouse was placed alone in the dark compartment and the mouse’s behavior was recorded during 5 minutes with a video camcorder located approximately 150 cm above the center of the box. Test was conducted during the morning. The latency before the first transition into the light compartment, the number of transitions between the two compartments and the time spent in each compartment were tested to assess for anxiety level and exploratory behavior, as published previously^92^.

### Olfactory preference test

This test is designed to identify specific detection deficiencies and/or odor preference, namely the ability to sense attractive or aversive scents. After 30 minutes of habituation to empty cage with no bedding, each mouse was challenged with a filter paper embedded with two strong scents (vanilla and 2-methyl butyrate) or a neutral scent (water) during 3 minutes and was video recorded. A one hour pause in between exposure to different scents was applied to each mouse using a procedure adapted from previously reported protocols^66,93^. The time the mouse spent sniffing the filter paper - the exploration time, is calculated *post hoc* by an examiner blind to mouse genotype and condition. Those scents with the exploration time greater than water were designated as “attractive” while those with times less than water were termed “aversive”.

### Social interaction in the home cage (resident-intruder test)

Social interaction was assessed in the home cage by a standard protocol^66,94^. Briefly, both resident and intruder mice were isolated and housed individually for 1 week before the task. After 30 minutes of habituation to the test room resident mouse was allowed to freely roam in his home cage without the cage top for 1 min. A novel male intruder mouse (non-littermate of same background, same age, and similar weight) was then introduced in the opposite corner as the resident and allowed to interact for 5 min while video-taped. The total physical interaction, defined as the time during which the resident mouse actively explores the intruder was analyzed *post hoc*. Only social activities, such as time spent investigating, grooming, following, sniffing etc. were quantified separately for each minute and for the whole time of the task and were differentiated from non-social/aggressive activities such as attacks, bites and tail rattles.

### Three-chamber social task

Specific social behaviors such as sociability and social recognition were analyzed by using three-chamber social task. The experimental procedure is adapted from Gascon E et al.^66^. The three-chamber box (59 × 39.5 × 21.5 cm) is made of transparent Plexiglas (Noldus Information Technology, Wageningen, The Netherlands) and is divided into three chambers (one middle and two side chambers) of equal size (18.5 × 39.5 cm) by the walls with a square opening (7 × 7 cm) that could be closed by a slide door. Each of the two side chambers contains a mobile wire cylinder shaped cage (20 × 10 cm diameter) that is made of transparent Plexiglas bars placed 6 mm apart. Cage is closed by the upper and lower lids.

Mice of both genotypes *(Fus*^+/+^ and *Fus*^*ΔNLS*/+^) that were experimentally tested are referred to as the test mice and adult male unfamiliar mice of same background, age and weight used as the social stimulus are called novel mice. All mice were housed individually for 1 week before the test and were habituated to the testing room for at least 1 hour before the start of behavioral tasks. One day prior to the testing, the novel mice were habituated to mobile wire cage for 5 min. The keeping of the novel mouse separated in a wire cage prevents aggressive and sexual activities, and in the same time ensures that any social interaction is initiated by the test mouse. Sessions were video-taped and visually analyzed *post hoc*. The experimental procedure was carried out in four trials of 5 min each. After each trial, the mouse was returned to his home cage for 15 min. Trials were grouped into two consecutive parts.

Trial 1 (habituation): the test mouse was placed in the middle chamber and left to freely explore each of the three chambers: the empty middle or two sides’ arenas containing the empty wire cages for 5 min.

Trials 2–4 (sociability, social recognition, social learning acquisition): the mouse was placed in the middle chamber, but an unfamiliar mouse (novel mouse) was placed into a wire cage in one of the side-chambers (the wire cage in the other side-chamber remains empty). The test mouse had free access to all three chambers. The position of novel mouse and empty wired cage were alternated between trials. We quantified the time spent actively exploring a novel mouse or an empty cage by the test mouse as a social interaction time or an object exploration time, respectively. The longer time that test mouse spents in the close perimeter around the cage containing the novel mouse while actively interacting with it (staring, sniffing) compared to the empty cage – object, indicates social preference or social recognition as a result of the capability to differentiate a conspecific from an object. The motivation of the test mouse to spontaneously interact with novel mouse is considered as sociability which gradually decreased over trials as a result of social learning acquisition..

### Water maze task

The water maze consisted of a circular pool (diameter 160 cm; height 60 cm) filled with water (21 ± 1 °C) made opaque by addition of a powdered milk (about 1.5 g/L). The habituation day consisted in one 4-trial session using a visible platform (diameter 11 cm, painted black, protruding 1 cm above the water surface and located in the South-East quadrant of the pool), starting randomly from each of the four cardinal points at the edge of the pool. During this habituation trial, a blue curtain surrounded the pool to prevent the use of distal cues and thus incidental encoding of spatial information. For the following days, the curtain was removed. Mice were given a 5-day training period (4 consecutive trials/day, maximum duration of a trial 60 seconds, inter-trial interval = 10–15 seconds) with a hidden platform located at a fixed position in the North-West quadrant. Animals were starting randomly from each of the four cardinal points at the edge of the pool and the sequence of the start points was randomized over days. Mice were tested for retention in a 18-days delay probe trial and two extinction tests: the first 2 hours after probe trial and the second 2 hours after the first. For the probe trial, the platform was removed; the mice were introduced in the pool from the North-East (a start point never used during acquisition) and allowed a 60seconds swimming time to explore the pool. Data were collected and computed by a video-tracking system (SMART; AnyMaze software). For the visible platform and training trials following parameters were used: the distance traveled and the latency time before reaching the platform and the average swimming speed. For the probe trial and extinction tests the time (in secondes) spent in the target quadrant (i.e., where the platform was located during acquisition) was analyzed^95^.

### Assessing neuronal activity by *in vivo* two-photon imaging

#### Cranial window implantation and virus injection

Mice of both sexes were implanted with a cranial window at 9 months of age (+/- 10 days) and received the injection of the genetically encoded calcium indicator (AAV2/1.Syn.GCaMP6m.WPRE.SV40 (diluted 1:6 in saline)^96^ into the primary motor cortex (M1), as described previously^97^. In brief, mice were first anesthetized with Fentanyl (0.05mg/kg), Midazolam (5.0mg/kg) and Metedomidin (0.5mg/kg). A circular craniotomy with a 2mm radius, centered 1.7mm lateral and 0.8mm anterior to bregma, was performed, followed by the slow injection of a total of ∼1ul of the calcium indicator into three sites (∼300nl per site at 600µm cortical depth). A 4mm round glass coverslip (Warner Instruments) was placed over the cortex and sealed with UV-curable dental acrylic (Venus Diamond Flow, Heraeus Kulzer GmbH). A metal head bar was attached to the skull with dental acrylic (Paladur, Heraeus Kulzer GmbH), allowing stable positioning for two-photon microscopy imaging.

#### Two-photon imaging in anesthetized mice

Four weeks following the cranial window implantation, *in vivo* two-photon imaging was performed within cortical layer 2/3 using a two-photon microscope (Hyperscope, Scientifica, equipped with an 8kHz resonant scanner) at frame rates of 30Hz and a resolution of 512×512 pixels. Using a 16x water-immersion objective (Nikon), stacks consisting of 15,000 frames (equivalent to ∼8 minutes) were acquired, covering a field of view (FOV) of 300×300 µm. Light source was a Ti:Sapphire laser with a DeepSee pre-chirp unit (Spectra Physics MaiTai eHP), as described previously ^97^. GCaMP6m was excited at 910nm, with a laser power not exceeding 40mW (typically 10 to 40mW). In each mouse, two or three FOVs at cortical depths of 140 to 310 µm were imaged, yielding 631 cells in *Fus*^+/+^ (n = 10 experiments, 5 mice) and 855 cells in *Fus*^ΔNLS/+^ mice (n = 14 experiments, 6 mice). During imaging, mice were anesthetized with 1.5% isoflurane in pure O_2_ at a flow rate of ∼0.5L/min, to maintain a respiratory rate in the range of 110 to 130 breaths per minute. Body temperature was maintained at 37 degrees using a physiological monitoring system (Harvard Apparatus).

#### Image processing and data analysis

All image analyses were performed in Matlab (Math Works) using custom-written routines ^97^. In brief, full frame images were corrected for potential x and y brain displacement, and regions of interests (ROIs) were semi-automatically selected based on the maximum and mean projections of all frames. Fluorescence signals of all pixels within a selected ROI were averaged, the intensity traces were low pass filtered at 10Hz. Contamination from neuropil signals was accounted for, as described previously^97^, using the following equation:

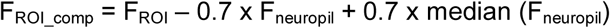

F_ROI_comp_ stands for neuropil-compensated fluorescence of the ROI, F_ROI_ and F_neuropil_ represent the initial fluorescence signal of the ROI and the signal from the neuropil, respectively. A neuron was defined as ‘active’ if it displayed at least one prominent calcium transient over 20 frames (corresponding to ∼0.7 seconds).

### Histological techniques

Mice were anesthetized with intraperitoneal injection of 100 mg/kg ketamine chlorhydrate (Imalgène 1000®, Merial) and 5mg/kg xylazine (Rompun 2%®, Bayer), and then transcardially perfused with cold PFA 4% in 0.01 M phosphate buffered saline (PBS). After dissection, brains were post-fixed for 24 hours and then included in agar 4% and serial cuts of 40 µm thick were made using vibratome (Leica Biosystems, S2000).

#### Peroxidase immunohistochemistry

For peroxidase immunohistochemistry, sections were incubated 10 min with H_2_O_2_ 3%, rinsed with PBS 1X and incubated with blocking solution (8% Horse serum, 0.3% Bovine Serum Albumin, 0.3% Triton, PBS-0.02% Thimerosal). Sections were rinsed, and then incubated with anti-mouse NeuN or anti-mouse parvalbumin (Millipore, MAB377, 1:100 and Sigma, P3088, 1:1000 respectively) overnight at room temperature. The second day, sections were rinsed and incubated for 2h at room temperature with biotinylated donkey anti-mouse antibody (Jackson, 715-067-003, 1:500). After sections were rinsed, they were incubated for 1h in horseradish peroxidase ABC kit (Vectastain ABC kit, PK-6100, Vector Laboratories Inc.), rinsed and incubated with DAB (Sigma, D5905). The enzymtic reaction was stopped by adding PBS 1X, rinsed with water and sections were mounted with DPX mounting medium (Sigma, O6522).

#### Quantification

Images were quantified using a homemade ImageJ plugin. A Region Of Interest (ROI) was first defined by the user as the M1/M2 regions of the cerebral cortex as defined by the Paxinos Atlas^98^ using the following coordinates: interaural 4.06 mm; Bregma 0.26 mm. For NeuN immunohistochemistry, a second, more anterior region of M1/M2 cortex was also quantified with the following coordinates: Interaural: 5.74mm, Bregma: 1.94mm.

In this region, a semi-automated segmentation led to the identification of the labelled structures (cells or nuclei). Finally, the plugin subdivided the previous ROI into 10 subregions and measured either the number of objects per subregion or the proportion of each subregion that is covered by labelled structures.

#### Immunofluorescence

Sections were rinsed with PBS 1X then incubated with blocking solution (8% Goat serum, 0.3% Bovine Serum Albumin, 0.3% Triton, PBS-0.02% Thimérosal) overnight at 4°C in primary antibody: rabbit anti-FUS antibody (ProteinTech, 11570-1-AP, 1:100) and mouse anti-parvalbumin antibody (Sigma, P3088, 1:1000). After 3 rinses in PBS, sections were incubated for 2h at room temperature with Hoechst (Sigma, B2261, 1/50.000) and secondary antibody: Goat anti-mouse Alexa-488 secondary antibody (Invitrogen, A11034, 1:500) and goat anti-mouse Alexa-647 secondary antibody (Invitrogen, A21245, 1:500). Finally sections were subsequently washed with PBS (3 x 10 min) and mounted in Aqua/polymount (Polysciences, 18606).

Images were acquired along the Z axis (Z stacking) using a Zeiss AxioImage.M2 microscope equipped with a Plan-Apochromat 20x/0.8 objective, high performance B/W camera (Orca Flash4, Hamamatsu) and run by the Zeiss Zen2 software. Images were quantified using the ImageJ freeware. First, the user defined ROIs corresponding to the cytoplasm and nucleus or several PV positive cells at several Z positions. Then a homemade macro was used to calculate the ratio, in the green channel, of the cytoplasm intensity divided by the nucleus one.

#### Electron microscopy

Mice were anesthetized by intraperitoneal injection of 100 mg/kg ketamine chlorhydrate and 5mg/kg xylazine and transcardiacally perfused with glutaraldehyde (2.5% in 0.1M cacodylate buffer at pH 7.4). Brains were dissected and immersed in the same fixative overnight. After 3 rinses in Cacodylate buffer (EMS, 11650), serial cuts of 80 µm thick were made with vibratome. Slides were then post-fixed in 1% osmium in Cacodylate buffer 1 hour at room temperature. Finally, tissues were dehydrated in graded ethanol series and embedded in Embed 812 (EMS, 13940). The ultrathin sections (50 nm) were cut with an ultramicrotome (Leica, EM UC7), counterstained with uranyl acetate (1% (w/v) in 50% ethanol) and observed with a Hitachi 7500 transmission electron microscope (Hitachi High Technologies Corporation, Tokyo, Japan) equipped with an AMT Hamamatsu digital camera (Hamamatsu Photonics, Hamamatsu City, Japan). Analysis of electron micrographs was performed as follows: 100 inhibitory synapses located in layers II/III were imaged per animal. Inhibitory synapses were identified as containing at least one mitochondria in each synaptic bouton. Synapses morphometry was analyzed using ImageJ freeware (National Institute of Health), where each synaptic boutons’ area was manually drawn as previously described^99^. An automated plugin was used to drawn and measure the active zones’ length, the number of synaptic vesicles within each bouton and the distance of each vesicle to the active zone, being as the beeline from the vesicle to the active zone. All images were acquired in layer II/III of the M1/M2 regions of the cerebral cortex as defined by the Paxinos Atlas^98^ using the following coordinates: interaural 4.06 mm; Bregma 0.26 mm.

### Synaptic density in brain sections

Mice were anesthetized by CO_2_ inhalation before perfusion with PBS containing 4% paraformaldehyde and 4% sucrose. Brains were harvested and post-fixed overnight in the same fixative and then stored at 4°C in PBS containing 30% sucrose. Sixty µm-thick coronal sections were cut on a cryostat and processed for free-floating immunofluorescence staining. Brain sections were incubated with the indicated primary antibodies (Rabbit GABAAalpha3 antibody Synaptic Systems, 1:500; Mouse Gephyrin antibody Synaptic Systems, 1:500, Guinea pig VGAT antibody, Synaptic Systems, 1/500) for 48 hours at 4°C followed by secondary antibodies (1:1000) for 24 hours at 4°C. The antibodies were diluted in 1X Tris Buffer Saline solution containing 10% donkey serum, 3% BSA, and 0.25% Triton-X100. Sections were then mounted on slides with Prolong Diamond (Life Technologies) before confocal microscopy.

Confocal images were acquired on a Leica SP8 Falcon microscope using 63X (NA 1.4) with a zoom power of 3. Images were acquired at a 2048×2048 pixel image resolution, yielding a pixel size of 30.05nm. To quantify the density of synaptic markers, images were acquired in the molecular layer 1/2 of the primary motor cortex area, using the same parameters for all genotypes. Images were acquired from top to bottom with a Z step size of 500nm. Images were deconvoluted using Huygens Professional software (Scientific Volume Imaging). Images were then analyzed as described^100^. Briefly, stacks were analyzed using the built-in particle analysis function in Fiji^101^. The size of the particles was defined according to previously published studies^72,102^. To assess the number of clusters, images were thresholded (same threshold per marker and experiment), and a binary mask was generated. A low size threshold of 0.01 um diameter and high pass threshold of 1 um diameter were applied. Top and bottom stacks were removed from the analysis to only keep the 40 middle stacks. For the analysis, the number of clusters per 40 z stacks was summed and normalized by the volume imaged (75153.8µm^3^). The density was normalized to the control group.

### Structural MRI scans

All data were acquired on a dedicated small bore animal scanner (Biospec 117/16, Bruker, Ettlingen, Germany) equipped with a cryogenically cooled two-element surface (MRI CryoProbeTM, Bruker BioSpec, Ettlingen, Germany) transmit/receive coil. Anatomical brain images were acquired in coronal slice orientation (30 slices) applying a gradient-echo (FLASH) sequence with acquisition parameters as: TE/TR 2.95/400 ms (TE = echo time, TR = repetition time), matrix 30 × 340 × 340, resolution 250 × 50 × 50 mm^3^). Volumetric tissue analysis followed previously established semi-automatic procedures^103,104^. Briefly, data processing was performed by the in-house developed software package Tissue Classification Software (TCS). For optimized visualization, the acquisition matrix of 340×340 voxels was transformed into a 768 × 768 grid by nearest neighbour interpolation. TCS includes mouse-based drawing tools for tissue/voxel selection. In order to define clearly visible regions, drawing was supported by a two-level thresholded conventional region-grow algorithm. Following the operator-defined intensity threshold, all connected voxels with respect to their intensity within the predefined intensity range were selected. The analysis was blinded and evaluated by the same experienced investigator (DW). Striatum, cortex, hippocampus, and ventricle sizes were identified in 30 slices. Volume sizes were normalized to intracranial volume (ICV), also determined by TCS.

### RNAseq

RNAseq on frontal cortex was performed as previously described^10,41^. Briefly, RNA from cortex of 22 months old *Fus*^ΔNLS/+^ mice and their control littermates were extracted with TRIzol (Invitrogen). RNA quality was measured using the Agilent Bioanalyzer system or RNA screenTape (Agilent technologies) according to the manufacturer’s recommendations. Samples were processed using the Illumina TruSeq single Stranded mRNA Sample Preparation Kit according to manufacturer’s protocol. Generated cDNA libraries were sequenced using an Illumina HiSeq 2000 sequencer with 4–5 biological replicates sequenced per condition using single read, 50 cycle runs. RNA from the cortex of 5 months old old *Fus* ^ΔNLS/+^ and control littermate mice was extracted and libraries were generated using the Illumina TruSeq single Stranded mRNA Sample Preparation Kit. The cDNA libraries were sequenced on a HiSeq 4000 with 3 biological replicates per condition using single-end 50bp read. Total reads sequenced varies from 35 to 45 million reads. Complete QC report will be made publicly available.

Raw sequencing files were processed using Salmon 1.2.0^105^ and mapped to the transcriptome. We generated a decoy-aware transcriptome using the entire mouse genome as a decoy sequence. The index was then created using the command as follow: - salmon index -t <FASTA> -I <INDEX> --decoys <DECOY> -k31.The quantification was then performed in —validateMappings mode as follow: - salmon quant -i <INDEX> -l <LIBTYPE> -r < FASTQ> --validateMappings -o <QUANT>. The tximport function from R3.6.2 was then used to import the quantification file. The counts and TPM values were then extracted using the counts and abundances tables respectively. Clustering and differential expression analyses. were performed with normalized counts. Differentially expressed genes were selected using pairwise comparisons with DESeq2 with a cut-off of *p*□<□0.1 after multiple testing correction. The enrichment tool EnrichR (*http://amp.pharm.mssm.edu/Enrichr)* v2016 was used for functional annotation of gene expression data^106^. Gene Ontology biological process and KEGG pathway enrichment analysis was conducted using the R tool for EnrichR with a threshold Benjamini-corrected *P*□≤□0.05. Common GO terms were identified by comparing different models and an enrichment score was calculated for each GO term based on the adjusted p-value and the number of genes detected. The Chord diagram was then plotted for the conserved GO terms with a custom Rscript which will be made available.

### Synaptosomal enrichment followed by RT-qPCR and western blotting

Frontal cortex was removed from the brains of 4 months old mice by micro-dissection, as previously described^107^, harvested, rapidly frozen in liquid nitrogen and stored at -80°C until use. Synaptosomal fraction was isolated using Syn-PER Synaptic Protein Extraction kit (Thermo Scientific, 87793) according to manufacturer’s instructions.

On synaptosomal preparations, RNA was extracted using TRIzol reagent (Sigma Aldrich, 93289). 1µg of RNA was reverse transcribed using iScript Ready-to-use cDNA supermix (Bio-Rad, 1708841). Quantitative PCR (qPCR) was performed using SsoAdvanced Universal SYBR Green Supermix (Bio-Rad, 172574) and quantified with Bio-Rad CFX Manager software. Gene expression was normalized by computing a normalization factor by Genorm software.using three standard genes *Pol2, Tbp* and *Actn* for nervous tissue. The following primer sequences were used:

*Pol2* : F- GCTGGGAGACATAGCACCA ; R- TTACTCCCCTGCATGGTCTC

*Tbp* : F- CCAATGACTCCTATGACCCCTA ; R- CAGCCAAGATTCACGGTAGAT

*Actn* : F- CCACCAGTTCGCCATGGAT ; R- GGCTTTGCACATGCCGGAG

*Fus* : F- TTATGGACAGACCCAAAAACACA ; R- TGCTGCCCATAAGAAGATTG

*Malat* : F- TAATGTAGGACAGCGGAGCC ; R- GTAGGGTAGTCCCCACTGCT

*Gad1* : F- CTGCGCCCTACAACGTATGA ; R- CACAGATCTTCAGGCCCAGTT

*Gad2* : F- CAGCTGGAACCACCGTGTAT ; R- TCAGTAACCCTCCACCCCAA

*Nrxn1* : F- ATGACATCCTTGTGGCCTCG ; R- TACTCTGGTGGGGTTGGCTA

*Chrnb2* : F- AACCCAGACGACTCCACCTA ; R- TGTAGAAGAGCGGTTTGCGA

*Chrna7* : F- CAGTGAGTGGAAGTTTGCGG ; R- GACATGAGGATGCCGATGGT

*Slc6a11* : F- AAGTGGTGCTGGAAAGTCGT ; R- AGCCCCAAGCAGGATATGTG

*Gabra1* : F- CCTCCCGAAGGTGGCTTATG ; R- CATCCCACGCATACCCTCTC

*Gabrb1* : F- TATGCTGGCGACATCGATCC ; R- AGTGGACCGACTGCTCAAAG

*Gabrb2* : F- TGGACCTAAGGCGGTATCCA ; R- GTGACTGCATTGTCATCGCC

*Gabrb3* : F- AGGAAGGCTTTTCGGCATCT ; R- ATGTTCCCGGGGTCGTTTAC

For western blotting cytosolic and synaptosomal fractions were prepared using the same protocol, and protein concentration was quantitated using the BCA protein assay kit (Pierce). 15µg of proteins were loaded into a gradient 4-20% SDS-PAGE gel (Bio-Rad, 5678094) and transferred on a 0.45µm nitrocellulose membrane (Bio-Rad) using a semi-dry Transblot Turbo system (Bio-Rad). Membranes were saturated with 10% non-fat milk in PBS and then probed with the following primary antibodies: Anti-Synaptophysin (Abcam, ab14692, 1:1000), Anti-FUS N-ter1 (ProteinTech, 11570, 1:1000), Anti-FUS N-ter2 (Bethyl, A-300-291A, 1/2000) and Anti-FUS C-ter (Bethyl, A300-294A, 1:2000) all diluted in 3% non-fat milk in PBS. Blot were washed and incubated with anti-Rabbit secondary antibody conjugated with HRP (P.A.R.I.S, BI2407, 1:5000) during 2 hours. Membranes were washed several times and analyzed with chemiluminescence using ECL Lumina Forte (Millipore, WBLUF0500) using the Chemidoc XRS Imager (Bio-Rad). Total proteins were detected with a stain-free gel capacity and normalized. Uncropped western blot images and stain-free images are provided in supplementary figures.

### Statistics

If not stated otherwise, data are presented as mean ± standard error of the mean (SEM). Statistical analyses were performed using GraphPad Prism 8 (GraphPad, CA). Unpaired t-test was used for comparison between two groups, one-way or two-way analysis of the variance (ANOVA), followed by Tukey’s multiple comparison *post hoc* test and two-way repeated measures (RM) ANOVA, followed by Sidak multiple comparison *post hoc* test were applied for three or more groups. Distributions were compared using the Kolmogorov Smirnov test. Results were considered significant when p<0.05. All statistical tests are reported in the **Table S1.**

## Figure legends

**Supplementary figure 1.**

(**a-c**) Bar charts showing time spent in peripheral and central quadrants of the open field arena of 10 months old *Fus*^+/+^ (black) and *Fus*^*ΔNLS*/+^ (orange) mice. Total distance traveled (in centimeters) (**middle graph**) and average speed of movements (in meters per seconds) (**left graph**) are presented. Data are presented as mean +/- SEM. N=6 per genotype. Multiple t-test. p=0.84 (peripheral quadrants) and p=0.34 (central quadrants). Unpaired t-test. p=0.13 (distance) and p=0.16 (velocity).

(**d**) Representative images of tracking’s trajectories (upper panels) and heat maps of mouse movement (lower panels) and in the open field of 10 months old *Fus*^+/+^ and *Fus*^*ΔNLS*/+^ animals.

(**e**) Bar charts showing latency time to enter the illuminated compartment (**left graph**), total time spent in the spent in the illuminated compartment (**middle graph**), and number of transitions between dark and the illuminated compartments (**right graph**) in the dark-light box test for 10 months old *Fus*^+/+^ (black) and *Fus*^*ΔNLS*/+^ (orange) mice. All values are represented as mean +/- SEM. N=14-15 per genotype. Unpaired t-test. p=0.15 (latency) and p=0.07 (total time) and p=0.11 (transitions).

**Supplementary figure 2.**

Analysis of olfactory function in *Fus*^+/+^ and *Fus*^*ΔNLS*/+^ mice at 22 months of age. Bar graphs showing similar preference of both genotypes for “attractive” scents (vanilla) compared to water and “aversive” stimuli. N=6 animals per genotype; All values are presented as mean +/- SEM of time mice spent sniffing filter paper immersed in corresponding solution. Multiple t-tests. p=0.74 (water), p=0.84 (vanilla) and p=0.64 (2-methyl butyrate).

**Supplementary figure 3.**

mRNA levels of the indicated genes in RNAs extracted from *Fus*^+/+^ (+/+) or *Fus*^ΔNLS/+^ (Δ/+) frontal cortex from 5 months old mice (Young) or 22 months old mice (Old) as assessed using RNAseq. All quantifications are presented relative to the +/+ Young levels set to 1. *, p<0.05 vs corresponding +/+ young levels, genome wide adjusted p-value.

**Supplementary Figure 4**

(**a**) Representative image of Parvalbumin immunohistochemistry of frontal cortex tissues at 10 and 22 months of age in *Fus*^+/+^ (+/+) or *Fus*^ΔNLS/+^ (Δ/+) mice.

(**b, c**) Distribution of parvalbumin+ neurons (PV+) in frontal cortex of Fus+/+ (black) vs FusΔNLS/+ (orange) at 10 (**b**) or 22 (**c**) months of age.

(**d**) Immunofluorescence of parvalbumin (red) and FUS (green) neuronal localization in frontal cortex in *Fus*^+/+^ (+/+) or *Fus*^ΔNLS/+^ (Δ/+) mice at 22 months of age.

(**e**) Violin plot showing the distribution of the ratio between nuclear and cytoplasmic in individual PV+ neurons in the frontal cortex of *Fus*^+/+^ (black) or *Fus*^ΔNLS/+^ (Δ/+) mice (orange) at 10 and 22 months.

**Supplementary Figure 5:**

Uncropped western blots and stain free labelling showing equal loading of gels of Figure 7.

