## Supplementary figures for "Cytoplasmic accumulation of FUS triggers early behavioral alterations linked to cortical neuronal hyperactivity and defects in inhibitory synapses"

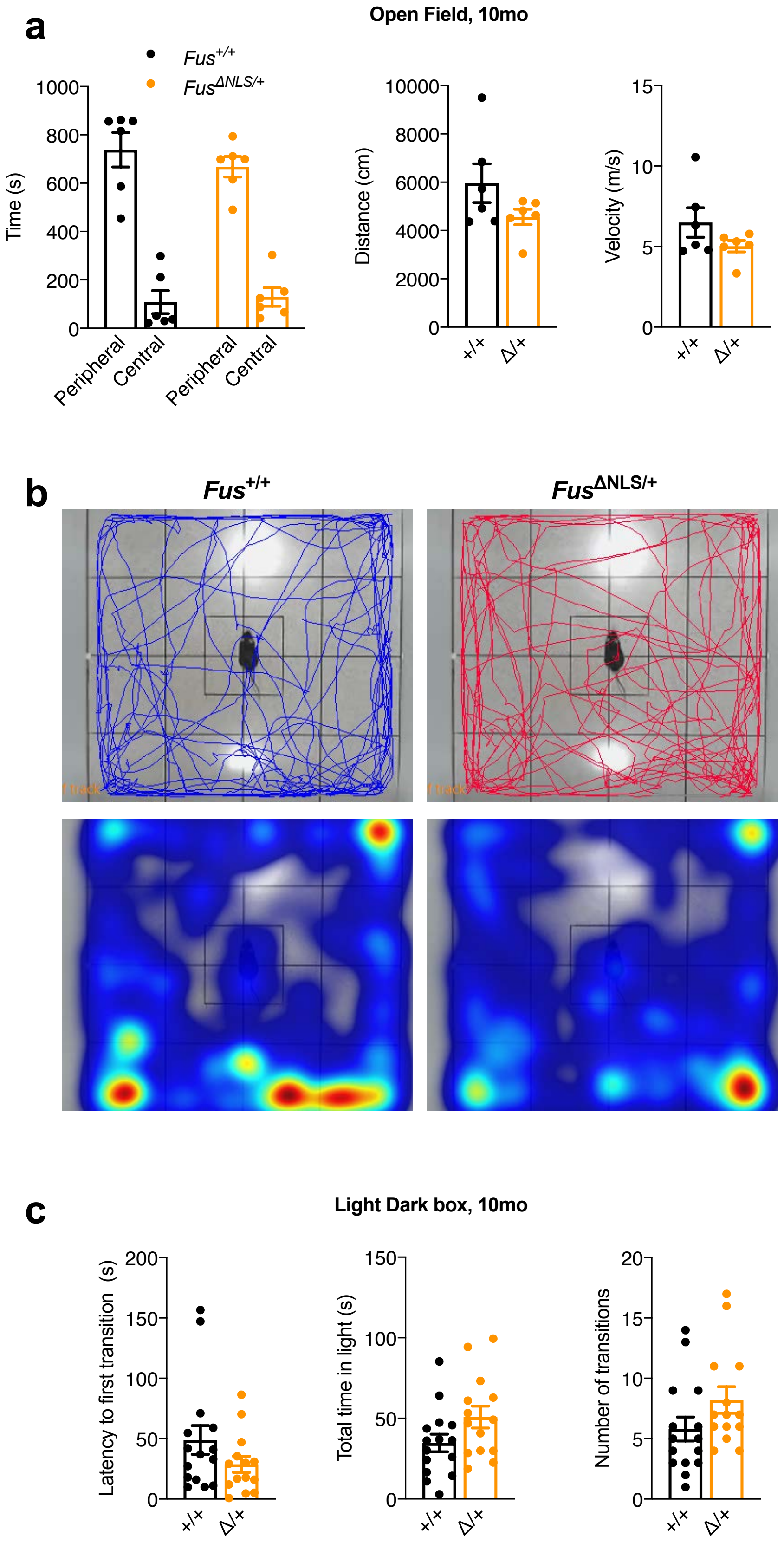

### Scekic-Zahirovic, Sanjuan-Ruiz et al, Figure S2

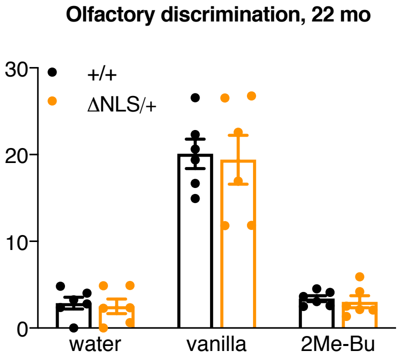

Scekic-Zahirovic, Sanjuan-Ruiz et al, Figure S3

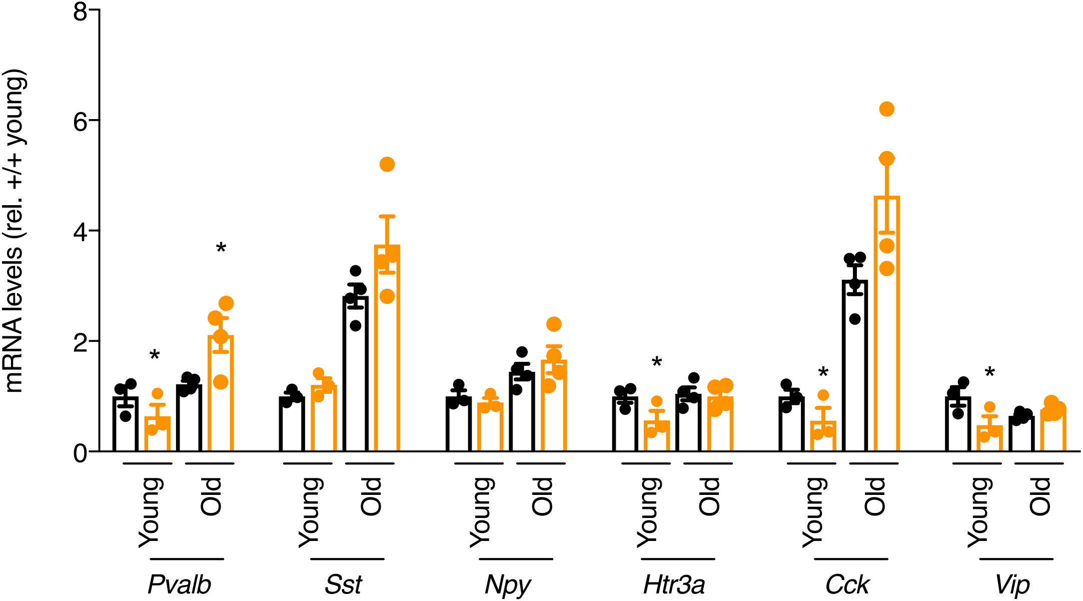

Scekic-Zahirovic, Sanjuan-Ruiz et al, Figure S4

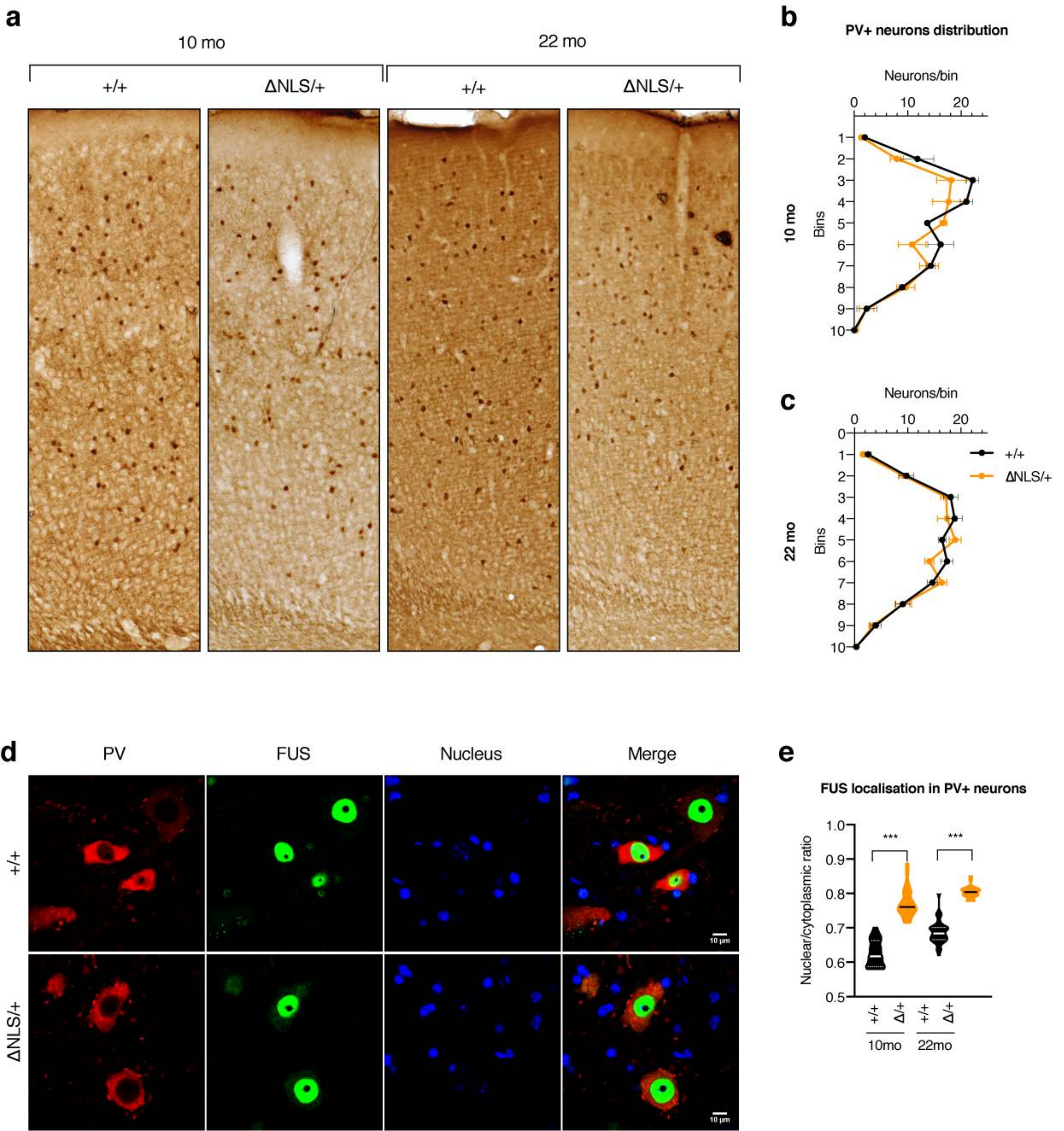

**Scekic-Zahirovic, Sanjuan-Ruiz et al., Figure S5**

FUS N-ter1

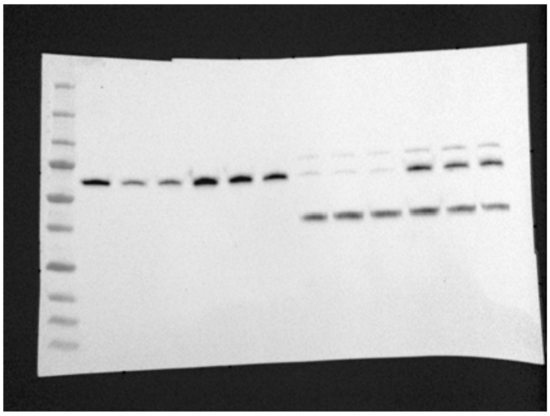

Stainfree for FUS N-ter1

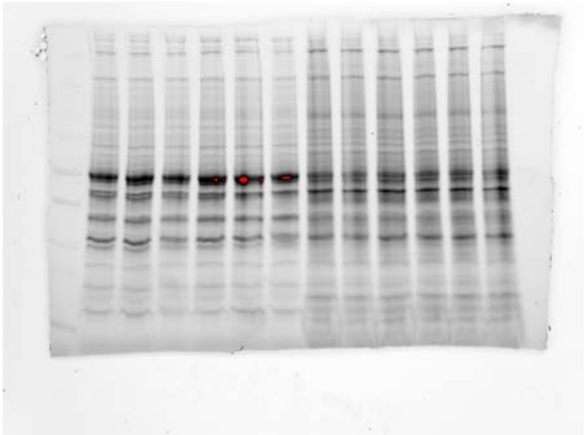

FUS N-ter2

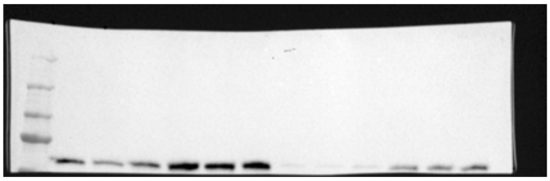

Synaptophysin

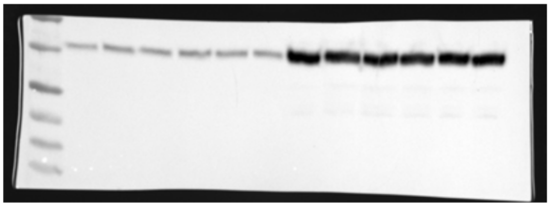

Stainfree for FUS N-ter 2
and synaptophysin

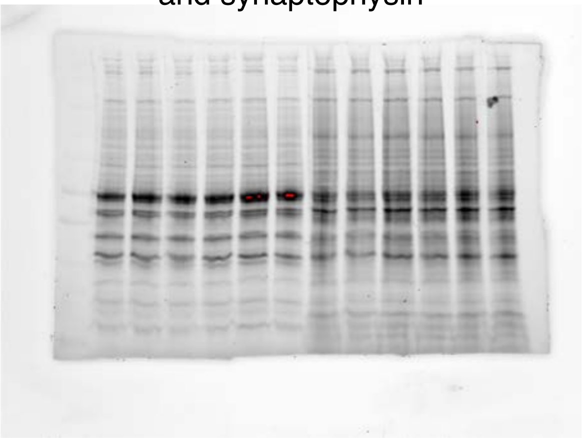

FUS C-ter

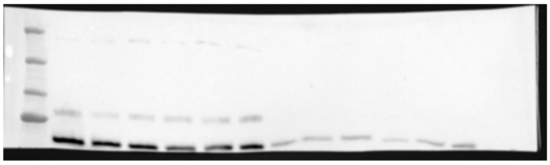

Stainfree for FUS C-ter

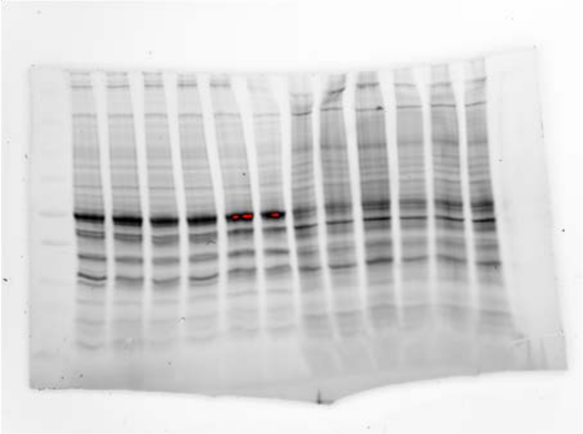
